# Desmoglein-1 coordinates cell proliferation and actin dynamics to promote epidermal morphogenesis

**DOI:** 10.64898/2026.09.29.755418

**Authors:** Brieanna L. Jarrell, Rosemary DiDominicis, Marihan Hegazy, Mariana De Niz, Robert Harmon, Xin Tong, Jennifer L. Koetsier, Dongmei Wang, Matthias Rübsam, Carien M. Niessen, Rui Yi, Lisa Godsel, Ziyou Ren, Kathleen J. Green

## Abstract

Skin development depends on coordinated biochemical and mechanical signaling to form a multilayered epidermal barrier. Our previous work demonstrated that the stratified tissue-specific desmosomal cadherin Desmoglein-1 (Dsg1) promotes epidermal differentiation in human *in vitro* models. Dsg1 first appears in the developing mouse epidermis around E14, coinciding with a developmental transition in stratification and the onset of terminal differentiation, raising the question of how Dsg1 contributes to epidermal morphogenesis *in vivo*. We generated a wild-type (WT) single-cell RNA sequencing (scRNA-seq) atlas of murine skin and integrated scRNA-seq data from Dsg1 knockout (Dsg1 KO) embryos spanning E14.5, E15.5, E16.5, and E18.5. Six keratinocyte subpopulations were annotated, including a Transitional subpopulation co-expressing basal and suprabasal markers, positioned between basal and differentiated states. Dsg1 KO epidermis exhibited reduced thickness at E14.5-E15.5, but recovered by E16.5-E18.5, suggesting compensatory adaptation. Differential gene expression analysis revealed enrichment of cell-cycle and actin-associated programs in Dsg1 KO during late development. EdU incorporation and PCNA staining confirmed increased proliferation in superficial layers accompanied by altered actin organization and changed cell morphology. Notably, actin and the Arp2/3 actin nucleation-promoting factor N-WASP exhibited marked changes in distribution during early and late development, respectively. Mechanistically, Dsg1 associates with Nck1, an adaptor linking N-WASP to Arp2/3-dependent actin polymerization. Silencing Nck1 or N-WASP reduced Dsg1-mediated stratification significantly, while a Nck1-binding-deficient Dsg1 mutant failed to rescue stratification in Dsg1-deficient keratinocytes *in vitro*. Together, these findings identify a Dsg1-Nck1-N-WASP axis linking desmosomal adhesion to cytoskeletal remodeling and support a model in which Dsg1 coordinates cell-cycle, cell morphology, and actin dynamics to drive epidermal morphogenesis.

## Introduction

Skin development requires the coordinated integration of mechanical and biochemical signaling to form a multilayered epidermis that provides a protective barrier capable of sensing and responding to environmental stresses throughout development and postnatal life^1^. Absolutely critical for barrier development and function are intercellular junctions, including gap junctions, tight junctions, adherens junctions (AJ), and desmosomes^2^. Collectively, these organelles are essential for communication across intracellular, cellular and tissue scales. In addition, they confer integrity to the tissue while contributing to dynamic morphogenetic mechanisms required for development.

Of these intercellular junctions, desmosomes are well-established to be critically important for contributing to tissue integrity through their ability to anchor tough and elastic keratin intermediate filaments to the plasma membrane via two cadherin subclasses, desmogleins and desmocollins^3^. The first immature desmosomes are seen at E8 in the single layer of surface ectoderm just prior to commitment to the epidermal keratinocyte lineage at E9.5^4^. This epidermal layer emerges from the surface ectoderm as a single layer of proliferative basal keratinocytes expressing Keratin-5/14 (Krt5/14), above which the periderm is formed around E11. At E12-13, basal keratinocytes transition from proliferating within a single layer to stratifying upward to form the developing epidermal layers. Multilayering is actively underway by E14, progressively giving rise to the spinous, granular, and terminally differentiated cornified layers that establish the epidermal barrier by E17.5 and persist after birth^5–7^.

Importantly, fully mature desmosomes form around E14.5 with the onset of intense multilayering and coinciding with the initial expression of one of the seven desmosomal cadherins, Desmoglein 1 (Dsg1)^8^. A terrestrial vertebrate innovation present exclusively in stratified epithelia, Dsg1 is first expressed in basal keratinocytes destined to differentiate and migrate to superficial layers^9^. Following progenitor cell commitment, Dsg1 forms a differentiation-dependent expression gradient with its highest expression in the superficial granular layers^8,10^. Consistent with its crucial role in epidermal integrity, loss of Dsg1 results in profound developmental defects resulting in perinatal lethality, as well as a spectrum of genetic autoimmune and pathogen-associated human disorders^8,11^. Despite the central importance of Dsg1 for epidermal development and barrier homeostasis, the extent to which it serves structural versus signaling roles *in vivo* during the dynamic developmental window spanning the onset of Dsg1 expression at E14.5 through mature barrier formation at E18.5 remains unknown.

Epidermal multilayering depends on expansion of the epidermis through proliferation as well as multiple stratification mechanisms. These include asymmetric cell divisions that generate basal and suprabasal daughter cells, and delamination events, in which differentiating basal cells move upward to the superficial layer^12^. Epidermal development is bi-phasic, with the initial early first phase occurring between E13.5-E15.5, where both basal and suprabasal keratinocytes have high proliferation rates and basal cells delaminate to fuel stratification. This is followed by a late second phase between E16.5-E18.5 where proliferation slows, particularly in the suprabasal layers, and delamination of basal cells may become a dominant mechanism supporting continued stratification^13^. Thus, a distinctive feature of early epidermal development is the capacity of stratified keratinocytes to continue dividing, a characteristic lost in late development and adult homeostasis^1,12–14^. Even so, how cell cycle processes in these stratified keratinocytes are developmentally regulated remains poorly understood.

While pathways driving cell cycle and proliferation during epidermal development are well-established, the mechanisms underlying mechanical changes important for epidermal keratinocyte delamination are less so. Studies of extrusion in simple epithelia and delamination in the epidermis have largely focused on alterations in contractile signaling pathways and junctional remodeling associated with classic cadherin-based junctions^15,16^. Recently, however, a role for desmosomes has begun to emerge in the process of apoptotic extrusion in simple epithelia, which predominantly express Dsg2 and Dsg3 rather than the stratified tissue-specific Dsg1^17,18^. We previously showed that ectopic expression of Dsg1 in simple epithelial MDCK cells is sufficient to drive delamination and formation of a second stratified layer, and Dsg1 is required in 3D organotypic cultures for delamination^10^. In human epidermal *in vitro* models, Dsg1 promotes differentiation, morphogenesis, and stratification through regulation of actin dynamics and tissue mechanics^19–22^. More specifically, we identified Arp2/3-dependent actin polymerization and consequent redistribution of cortical tension as potential drivers of Dsg1-dependent delamination^10^. Nevertheless, the extent to which Dsg1 contributes to the process of stratification *in vivo* is not well-characterized.

Single-cell RNA sequencing (scRNA-seq) has advanced our understanding of skin development by revealing keratinocyte heterogeneity and potential drivers of epidermal stratification during early murine development (E12.5-E14.5) prior to barrier formation at E16.5^23,24^. However, a developmental atlas spanning the transition from the early, highly proliferative phase to later stages of stratification and terminal differentiation is lacking. To bridge this gap, we generated a wild-type (WT) murine skin scRNA-seq atlas spanning E14.5, E15.5, E16.5, and E18.5, enabling us to define cellular and molecular changes accompanying this developmental transition. Given that Dsg1 integrates epidermal differentiation, barrier function and stratification through delamination in human *in vitro* systems^10,19,20^, we extended this analysis to Dsg1 KO embryos and complemented these *in vivo* studies with human mechanistic models to determine how Dsg1 contributes to epidermal morphogenesis.

Comparison of Dsg1 KO and WT epidermis across this developmental window revealed an early defect in tissue thickness followed by later recovery, which is associated with changes in proliferation, cell morphology and actin organization. These observations pointed to cytoskeletal remodeling as a potential mechanism through which Dsg1 regulates epidermal morphogenesis. Consistent with this possibility, actin and the Arp2/3 actin nucleation-promoting factor, N-WASP, exhibited profound alterations in distribution in Dsg1 KO epidermis. We further identified a predicted Nck1 SH2-binding phosphotyrosine motif at Y1042 in the Dsg1 cytoplasmic tail and validated Nck1 as a functional link between Dsg1 and the N-WASP:Arp2/3 actin remodeling machinery. Nck1 association was required for Dsg1-dependent stratification *in vitro*, identifying a Dsg1-Nck1-N-WASP axis through which desmosomal adhesion can engage actin remodeling to promote proper epidermal morphogenesis.

## Results

### Subclustering of Dsg1 WT and Dsg1 KO keratinocytes identifies a Transitional subpopulation co-expressing basal and suprabasal markers

To define keratinocyte heterogeneity across epidermal development and determine how loss of Dsg1 affects this process, we performed scRNA-seq on dorsal skin from Dsg1 WT and Dsg1 KO embryos spanning E14.5, E15.5, E16.5, and E18.5 (**Figure 1A**). We first generated a Dsg1 WT developmental atlas comprising 39,102 cells, encompassing the transition from early epidermal multilayering through mature barrier formation (**Figure S1A**) and subsequently used this atlas as a reference to annotate 62,459 Dsg1 KO cells from matched developmental timepoints (**Figure 1A**). To integrate the datasets and mitigate batch effects, Harmony was employed, aligning cellular heterogeneity across different developmental stages^25^. Subsequently, Seurat clustering classified cells based on gene expression profiles, allowing the identification of eight general cell populations by utilizing the Seurat v5 FindMarkers function across all timepoints^26^ (**Figure S1B**).

**Figure 1:**
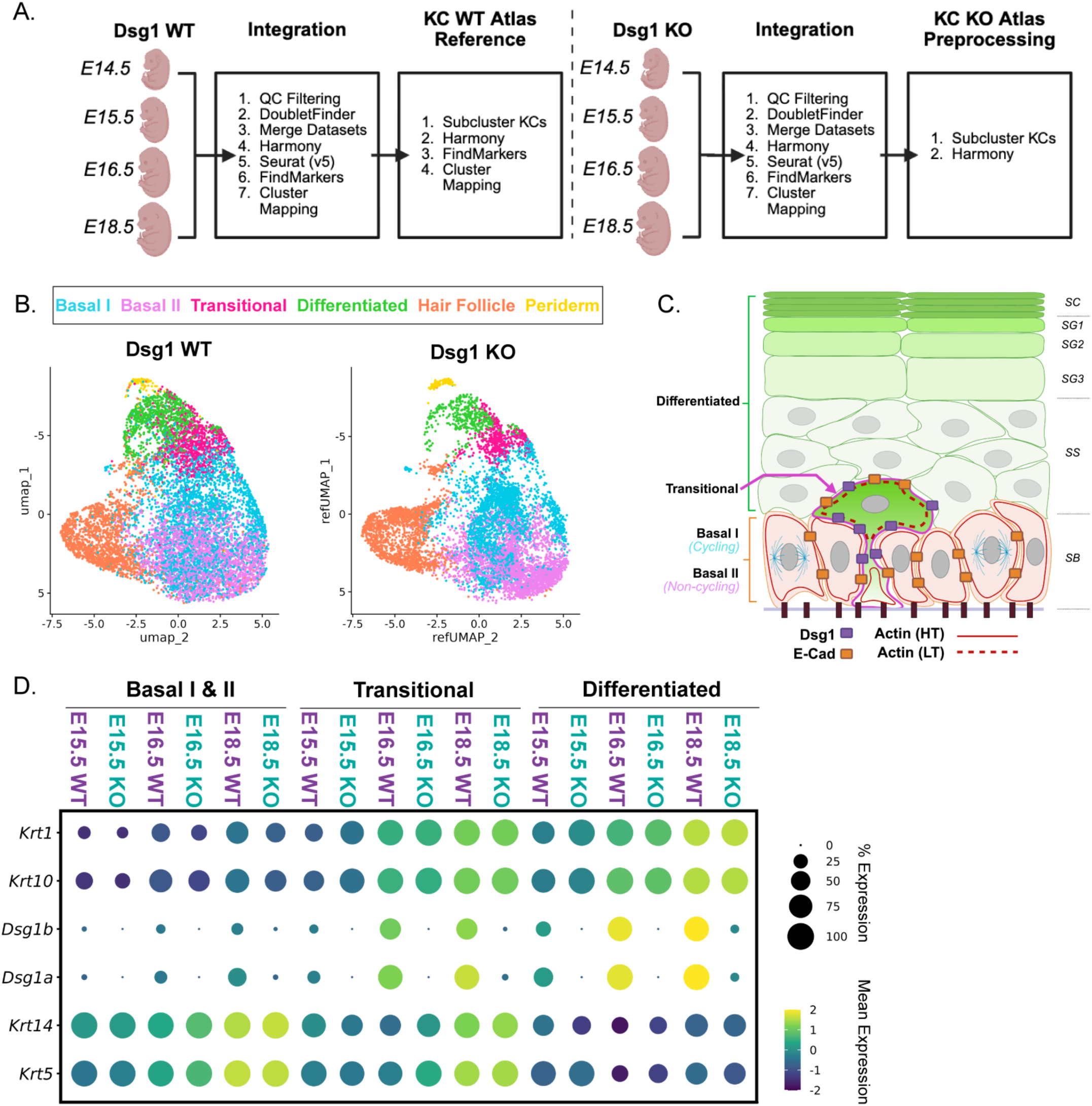
Dsg1 WT and Dsg1 KO scRNA-seq integration defines six keratinocyte subpopulations across epidermal development, including a uniquely resolved Krt5/14+ and Krt1/10+ expressing Transitional subpopulation. A) Workflow schematic of the scRNA-seq integration pipeline for generation of the Dsg1 WT reference atlas and preprocessing of the Dsg1 KO dataset across developmental timepoints (E14.5, E15.5, E16.5 & E18.5) for downstream analyses. B) Combined developmental timepoint UMAPs of the WT reference (left panel) and KO query (right panel) showing all six keratinocyte subpopulations: Basal I (cycling), Basal II (non-cycling), Transitional, Differentiated, Hair Follicle, and Periderm. Keratinocyte subpopulations are color coded. C) Epidermal cartoon schematic highlighting the four keratinocyte subpopulations selected for downstream analyses: Basal I, Basal II, Transitional, and Differentiated. D) Dot plot of selected genes across E15.5, E16.5, & E18.5 in combined Basal I & II, Transitional, and Differentiated subpopulations, comparing Dsg1 WT and Dsg1 KO. Note increased expression of Krt5/14, Krt1/10, and Dsg1a/b in the Transitional subpopulation across all Dsg1 WT timepoints.

To further investigate keratinocyte-specific changes across epidermal development, we subclustered the keratinocyte population and resolved six distinct subpopulations (**Figure S2A-B**). These included well-characterized populations such as Hair Follicle keratinocytes, the early protective Periderm, and suprabasal Differentiated keratinocytes, as well as three basal progenitor subpopulations: proliferative Basal I, quiescent Basal II, and Transitional keratinocytes (**Figure 1B-C, S2D**). Notably, Transitional keratinocytes co-expressed the basal markers, keratin 5 and 14 (Krt5/14), suprabasal markers, keratin 1 and 10 (Krt1/10) and Dsg1, indicating the initiation of differentiation and supporting their identity as a potential intermediate state between Basal I/II and Differentiated keratinocyte subpopulations (**Figure 1D**).

To determine how loss of Dsg1 alters these keratinocyte populations across development, we used the Dsg1 WT murine skin developmental atlas as an anchor reference to annotate Dsg1 KO cells using the Seurat label-transfer workflow (**Figure 1B, S2A,C)**. This approach enabled direct comparison of corresponding keratinocyte subpopulations between Dsg1 WT and Dsg1 KO embryos across early (E14.5 and E15.5) and late (E16.5 and E18.5) developmental stages. The integrated dataset was subsequently used for differential gene expression and gene enrichment analysis^27^, as well as Palantir pseudotime analysis^28^, to investigate Dsg1-dependent changes in transcriptional programs, cell-state transitions, and developmental trajectories (**Figure 2A-C, S2D**). Together, this framework allowed us to examine how loss of Dsg1 alters keratinocyte states and developmental progression.

**Figure 2:**
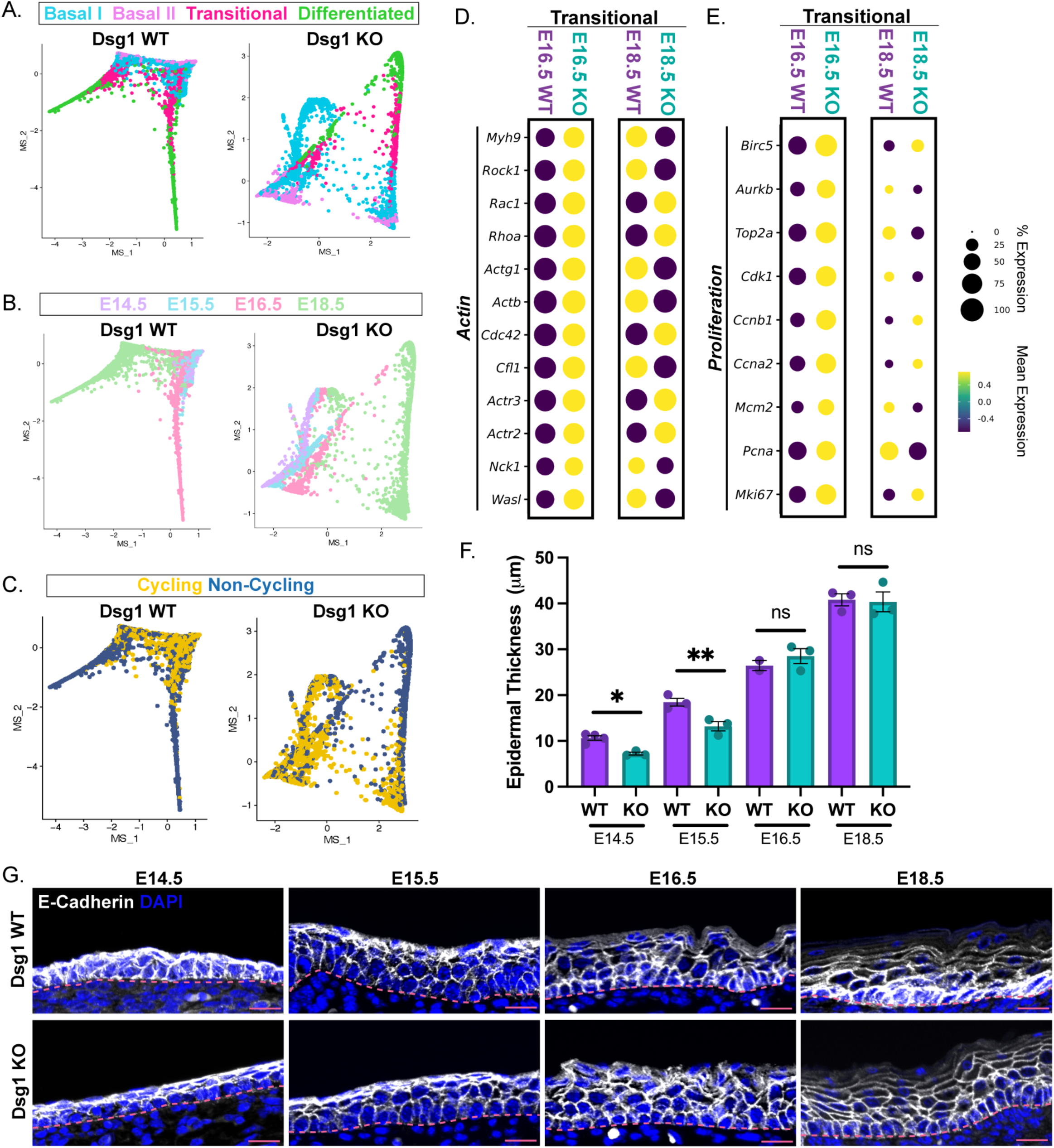
Transitional keratinocytes define an intermediate state between Basal I/Il and Differentiated transcriptional cell states, with altered cell cycle and actin-associated gene programs accompanying an early epidermal thickness defect that recovers during late development in Dsg1 KO epidermis. A) Palantir pseudotime trajectory across four keratinocyte subpopulations: Basal II, Basal I, Transitional, and Differentiated. B) Palantir pseudotime trajectory colored by developmental timepoint (E14.5, E15.5, E16.5, & E18.5). C) Palantir pseudotime trajectory colored by cell cycle state (Cycling & Non-Cycling). D) Dot plot of selected actin-associated genes in the Transitional subpopulation at E16.5 & E18.5, comparing Dsg1 WT and Dsg1 KO. Note the increased expression of actin-associated genes in Dsg1 KO compared with Dsg1 WT. E) Dot plot of selected cell proliferation-associated genes in the Transitional subpopulation at E16.5 & E18.5, comparing Dsg1 WT and Dsg1 KO. Note the increased expression of cell proliferation-associated genes in Dsg1 KO compared with Dsg1 WT. F) Quantification of epidermal thickness from H&E-stained sections comparing Dsg1 WT and Dsg1 KO across developmental timepoints E14.5, E15.5, E16.5, & E18.5. (mean ± SEM; n=2-4 embryos per genotype/timepoint; unpaired two-tailed Welch’s t-test) G) Representative images of E-cadherin-stained dorsal skin tissue sections from Dsg1 WT and Dsg1 KO across developmental timepoints E14.5, E15.5, E16.5, & E18.5. The dotted line marks the basement membrane. Scale bar 20µm. p-values are represented by (*) = 0.05, (**) = 0.01; ns = not significant.

### The loss of Dsg1 disrupts cell fate, temporal trajectory, and gene expression profiles across early and late development

Single-cell differentiation trajectories were used to determine whether Transitional keratinocytes occupy the proposed intermediate state between Basal I/II and Differentiated keratinocyte populations. This analysis was restricted to the Dsg1 WT interfollicular epidermal subset, comprising Basal I, Basal II, Transitional, and Differentiated keratinocytes (**Figure 1C**). Palantir multiscale space was used to provide a low-dimensional representation to preserve local and global cellular relationships to visualize differentiation trajectories^28^. Cell identities previously defined by gene expression in Seurat were mapped onto the multiscale space (**Figure 2A**), followed by developmental timepoint (**Figure 2B**) and cell-cycle state (**Figure 2C**) to examine their distribution along the inferred cell state trajectory. The trajectory analysis of Dsg1 WT revealed that the Transitional keratinocytes occupy the proposed intermediate position (**Figure 2A, left panel**). Intriguingly, applying this analysis to the Dsg1 KO dataset revealed that the Dsg1 KO displays diffuse cell fate trajectory organization, suggesting that the loss of Dsg1 alters normal cell fate trajectories (**Figure 2A, right panel**). Furthermore, trajectory analysis of Dsg1 WT dataset was relatively linear and temporally ordered trajectory, whereas the Dsg1 KO displayed a more diffuse trajectory with reduced separation between developmental timepoints, particularly between E16.5 and E18.5 (**Figure 2B**).

Given the central role of Dsg1 in desmosomal adhesion and its emergence during epidermal stratification, we next asked whether Dsg1 contributes to the organization of broader junctional and cytoskeletal programs during development. Gene signature analysis revealed alterations in several desmosomal, intermediate filament, select tight junctional, actin and cell proliferation-associated genes in Dsg1 KO, whereas no notable changes were observed among the selected AJ-associated genes among Basal I/II, Transitional, and Differentiated subpopulations (**Figure S3A-C, S4A-B**). In the Dsg1 WT dataset, *Dsg1a/b* expression progressively increased from Basal I/II to Transitional to Differentiated subpopulations across E15.5, E16.5 & E18.5, consistent with Dsg1’s association with epidermal maturation (**Figure S3A**). Extending these observations, trajectory analysis revealed elevated cell cycling gene expression in Dsg1 KO cells, compared with Dsg1 WT cells (**Figure 2C**), while analysis of the Transitional subpopulation during late development (E16.5, E18.5) showed enrichment of both actin-associated and cell proliferation signatures in the Dsg1 KO (**Figure 2D-E**).

We next asked whether these transcriptional signatures were accompanied by alterations in epidermal architecture. Histological analysis of H&E-stained dorsal skin together with E-cadherin, a classical AJ marker used to delineate the living layers of the epidermis, revealed that Dsg1 KO epidermis was significantly thinner than Dsg1 WT at E14.5 and E15.5. This defect began to resolve by E16.5, with epidermal thickness restored to Dsg1 WT levels by E18.5 (**Figure 2F**) while E-cadherin distribution remained comparable between genotypes throughout development (**Figure 2G)**. Together, these findings suggest that loss of Dsg1 disrupts epidermal stratification, followed by potential adaptive or compensatory mechanisms that restore epidermal thickness prior to birth.

### The loss of Dsg1 promotes the mislocalization and elevation of cell proliferation during late development

Loss of Dsg1 disrupts the suppression of EGFR-ERK1/2 and MAPK signaling normally associated with commitment to epidermal differentiation^19^. Together with the increased proliferation-associated gene signature identified in our scRNA-seq analysis of Dsg1 KO keratinocytes, this prompted us to ask whether loss of Dsg1 alters the cell-cycle dynamics during epidermal development. 1-hr EdU incorporation revealed no difference between Dsg1 WT and Dsg1 KO epidermis at E15.5, but a significant increase emerged by E16.5 and persisted at E18.5 (**Figure S5A-B**). Consistent with these findings, PCNA+ cells were also increased in Dsg1 KO epidermis at E16.5 and E18.5, but not E15.5 (**Figure 3A**). To determine the spatial distribution of this increased proliferation, we performed concurrent Krt14, Krt10, and PCNA immunofluorescence. PCNA+/Krt10+ suprabasal and PCNA+/Krt10+/Krt14+ transitional populations were significantly increased in the Dsg1 KO epidermis (**Figure 3C-E**), whereas the proportion of PCNA+/Krt14+ basal cells was unchanged (**Figure 3B,E**). These findings suggest that the late developmental increase in proliferation is driven primarily by persistent ectopic cell-cycle activity in suprabasal compartments in the absence of Dsg1.

**Figure 3:**
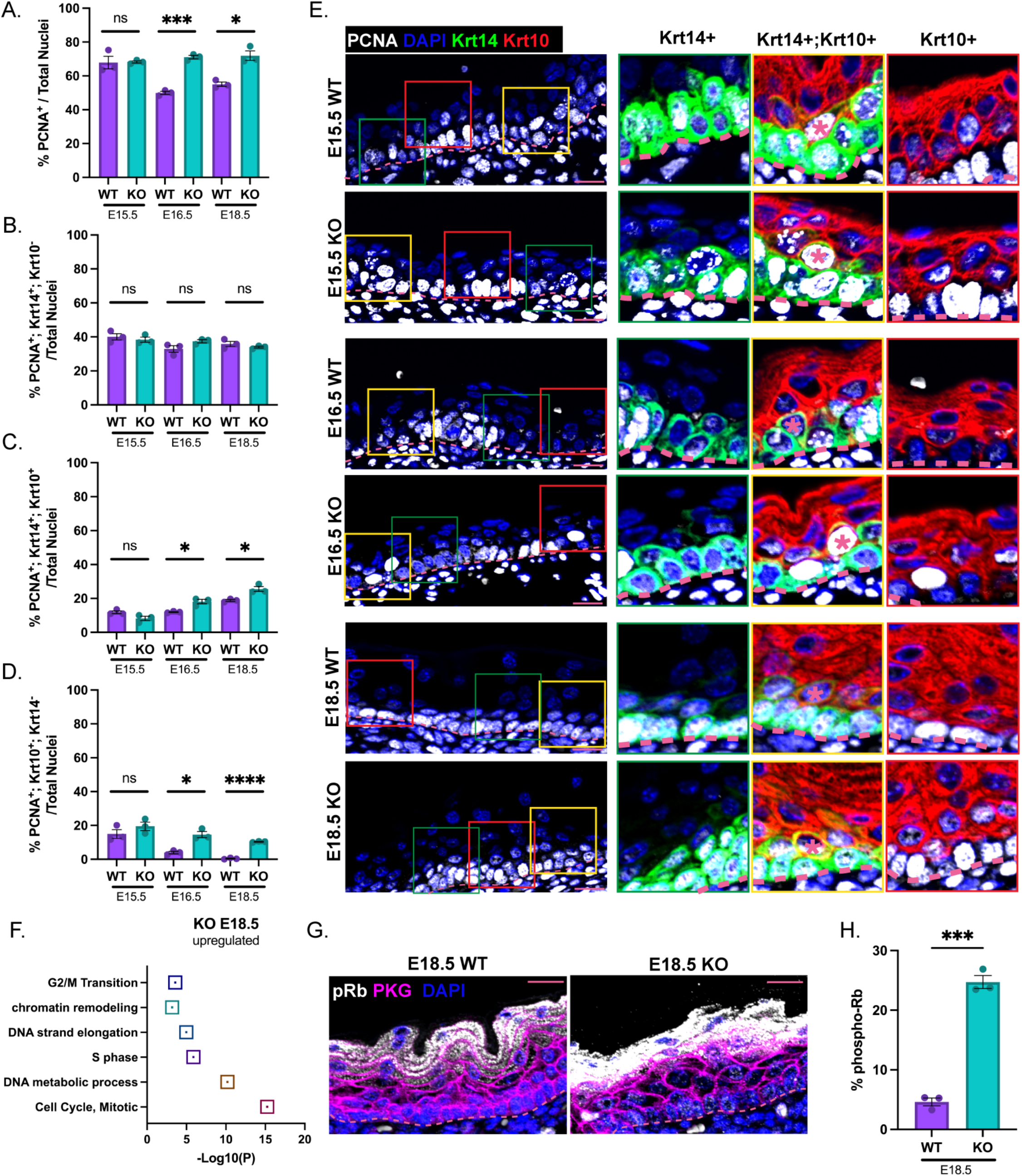
Atypical cell-cycle activity occurs in the transitional and differentiated keratinocytes during late development in Dsg1 KO epidermis. A) Quantification of the percentage of PCNA-positive cells across developmental timepoints E15.5, E16.5, & E18.5, comparing Dsg1 WT and Dsg1 KO. B) Quantification of Keratin-14/PCNA double-positive basal cells shown in (E, first zoom inset, Krt14+). C) Quantification of Keratin-14/Keratin-10/PCNA triple-positive transitional cells, shown in (E, middle zoom inset, Krt14+;Krt10+, denoted by pink asterisk). D) Quantification of Keratin-10/PCNA double-positive suprabasal cells shown in (E, last zoom inset, Krt10+). E) Representative images of dorsal skin tissue sections from Dsg1 WT and Dsg1 KO at E15.5, E16.5, and E18.5 stained for Keratin-14, Keratin-10, PCNA, and DAPI. F) Metascape Pathway enrichment analysis showing E18.5 Dsg1 KO total keratinocyte upregulation of cell-cycle related pathways. G) Representative images of dorsal skin tissue sections from E18.5 Dsg1 WT and Dsg1 KO stained for pRb, plakoglobin (PKG), and DAPI. I) Quantification of the masked percentage of total pRb signal shown in (G), excluding staining within the uppermost stratum granulosum and stratum corneum layers. The dotted line marks the basement membrane. Data are presented as mean ± SEM; n = 3 embryos per genotype/timepoint; unpaired two-tailed Welch’s t-test. Scale bar 20µm. p-values are represented by (*) = 0.05, (**) = 0.01, (***) = 0.001; ns = not significant.

To define signaling changes that could support continued cell cycling, we performed Metascape enrichment analysis on genes upregulated in E18.5 Dsg1 KO keratinocytes, which revealed enrichment of S phase, G2/M transition and broader cell cycle-related pathways (**Figure 3F**). Because cell-cycle progression is regulated by the Rb-E2F signaling axis, in which mitogen-induced Rb phosphorylation relieves E2F repression to promote progression of cell-cycle gene expression^29,30^, we examined Rb phosphorylation at E18.5. Dsg1 KO epidermis exhibited increased phospho-Rb (pRb)-positive nuclei (**Figure 3G-H**), consistent with enhanced cell-cycle progression. Together with the increased EdU incorporation and expansion of PCNA+/Krt14+/Krt10+ transitional and PCNA/Krt10+ suprabasal populations, these findings indicate that loss of Dsg1 permits persistent cell-cycle activity during late epidermal development.

### Dsg1 loss alters the temporal and spatial morphogenetic program

Cell-cycle exit is closely coupled to the morphologic remodeling that accompanies epidermal stratification, as proliferative basal keratinocytes transition from small, cuboidal shapes into rounded morphologies during mitosis, and subsequently shift into larger, polyhedral or flattened shapes as basal cells delaminate and move upward into the suprabasal spinous layer^31,32^. Our previous *in vitro* studies showed that Dsg1 promotes differentiation in human models by attenuating Ras-Raf-ERK pathway signaling without inhibiting basal cell proliferation^19,20^. We therefore asked whether the persistence of proliferative activity observed in Dsg1 KO epidermis was accompanied by changes in cellular architecture and tissue organization.

Loss of Dsg1 produced progressive, compartment-specific changes in keratinocyte morphology. At E15.5 whole-mount imaging of skin stained with phalloidin revealed an increase in the volume of Krt14+ basal cells in the Dsg1 KO epidermis, while Krt10+ suprabasal cell volume was unchanged (**Figure 4A-B, S6A-B**). By E16.5 increased cell volume extended to both Krt14+ basal and Krt10+ suprabasal compartments, coinciding with recovery of epidermal thickness (**Figure 4C-D, S7A-B**).

**Figure 4:**
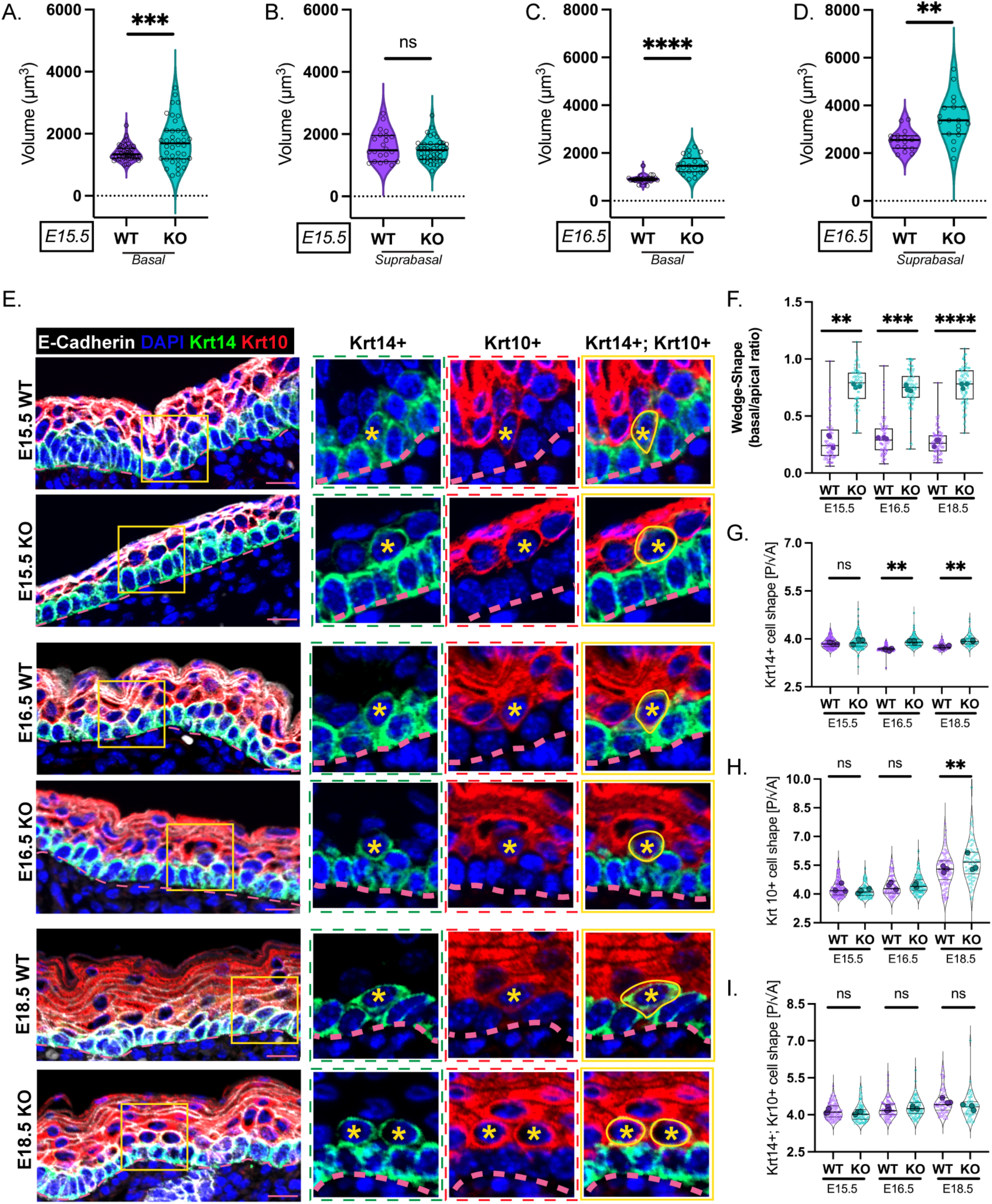
Loss of Dsg1 alters cell shape morphology. A) Quantification of basal keratinocyte cell volume from E15.5 dorsal skin whole-mount images stained with phalloidin, comparing Dsg1 WT and Dsg1 KO. B) Quantification of suprabasal keratinocyte cell volume from E15.5 dorsal skin whole-mount images stained with phalloidin, comparing Dsg1 WT and Dsg1 KO. C) Quantification of basal (Krt14+) keratinocyte cell volume from E16.5 dorsal skin whole-mount images stained with phalloidin, comparing Dsg1 WT and Dsg1 KO. D) Quantification of suprabasal keratinocyte cell volume from E16.5 dorsal skin whole-mount images stained with phalloidin, comparing Dsg1 WT and Dsg1 KO. E) Representative images of dorsal skin tissue sections from Dsg1 WT and Dsg1 KO at E15.5, E16.5, & E18.5 stained for E-cadherin, Keratin-14 (Krt14+), Keratin-10 (Krt10+), and DAPI. Yellow asterisks indicate Krt14+/Krt10+ double-positive cells, with corresponding yellow outlines highlighting their wedge-like or round-like cell morphology. The dotted line marks the basement membrane. F) Quantification of the basal-to-apical width ratio as a measure of wedge-like cell morphology in Krt14+/Krt10+ double-positive cells shown in (E, last zoom inset, Krt14+;Krt10+). G-I) Quantification of cell shape index for Krt14+ single-positive basal cells, Krt14+/Krt10+ double-positive transitional cells, and Krt10+ single-positive suprabasal cells, respectively, comparing Dsg1 WT and Dsg1 KO across E15.5, E16.5, and E18.5. (A-D) Individual data points represent single-cell measurements pooled from n = 2-3 embryos per genotype/timepoint; cell-level distributions were performed using the two-sample Kolmogorov-Smirnov test. (F-I) Individual cell measurements are shown as smaller circles, with larger circles representing the mean for each individual embryo; n = 3 embryos per genotype/timepoint. Statistical comparisons were performed on embryo-level means using an unpaired two-tailed Welch’s t-test. Scale bar 20µm. p-values are represented by (**) = 0.01, (***) = 0.001, (****) = 0.0001; ns = not significant.

Because basal-to-suprabasal transition requires directional changes in cell geometry, we next examined transitional Krt14+/Krt10+ keratinocytes. Given the established role of Dsg1 in promoting keratinocyte delamination, these intermediate cells provide a potential link between Dsg1-dependent changes in cell morphology and the transition from basal to suprabasal cell states. In Dsg1 WT epidermis stained for E-cadherin, these cells exhibited a characteristic wedge-like morphology, whereas Dsg1 KO transitional cells were more rounded (**Figure 4E**), reflected by an altered basal-to-apical width ratio (**Figure 4F**). We then asked whether broader changes in cell shape accompanied this altered transitional geometry by quantifying the cell shape index (q = perimeter [P]/ √ area [A])^33,34^ across basal (Krt14+), transitional (Krt14+/Krt10+), and suprabasal (Krt10+) compartments (**Figure 4G-I)**. Although shape index was similar between genotypes at E15.5, Dsg1 KO basal cells showed an increased shape index at E16.5 and E18.5 (**Figure 4G)**. No significant differences were observed in transitional Krt14+/Krt10+ double positive cells across the three developmental timepoints (**Figure 4H**), whereas Dsg1 KO suprabasal (Krt10+) keratinocytes exhibited an increased cell shape index at E18.5 (**Figure 4I)**. Together, these findings reveal a progression from temporal alterations in basal cell volume and apical-to-basal geometrical asymmetry in Krt14+/Krt10+ double positive cells to broader, compartment-specific changes in cell shape during later epidermal development, highlighting the role of Dsg1 in regulating distinct aspects of keratinocyte geometry throughout development.

### The loss of Dsg1 disrupts actin organization and localization of the actin regulator and Arp2/3 nucleation-promoting factor, N-WASP

Because cell-shape remodeling requires dynamic reorganization of the actin cytoskeleton, we next asked whether the morphogenetic defects observed in Dsg1 KO epidermis were associated with altered actin organization. Phalloidin whole-mount imaging at E15.5 and E16.5 was used to calculate actin coherency, a measure of local actin alignment^35–37^. At E15.5, actin coherency was significantly higher in both basal (Krt14+) and suprabasal (Krt10+) compartments of Dsg1 WT compared with Dsg1 KO epidermis (**Figure 5A,C-E, S6A-B**). Dsg1 WT basal cells also exhibited greater actin enrichment at tricellular junctions (TCJ) relative to bicellular junctions (BCJs), which was reduced in Dsg1 KO basal cells (**Figure 5B**). Notably, Dsg1 KO basal cells exhibited increased actin enrichment at the midpoint of BCJs, suggesting altered junctional actin organization (**Figure 5A**). Reduced actin coherency persisted in both basal and suprabasal compartments in Dsg1 KO at E16.5 (**Figure S7A-B, Figure S8A-D**). These data suggest that the loss of Dsg1 disrupts the organization and junctional distribution of the actin cytoskeleton during epidermal development.

**Figure 5:**
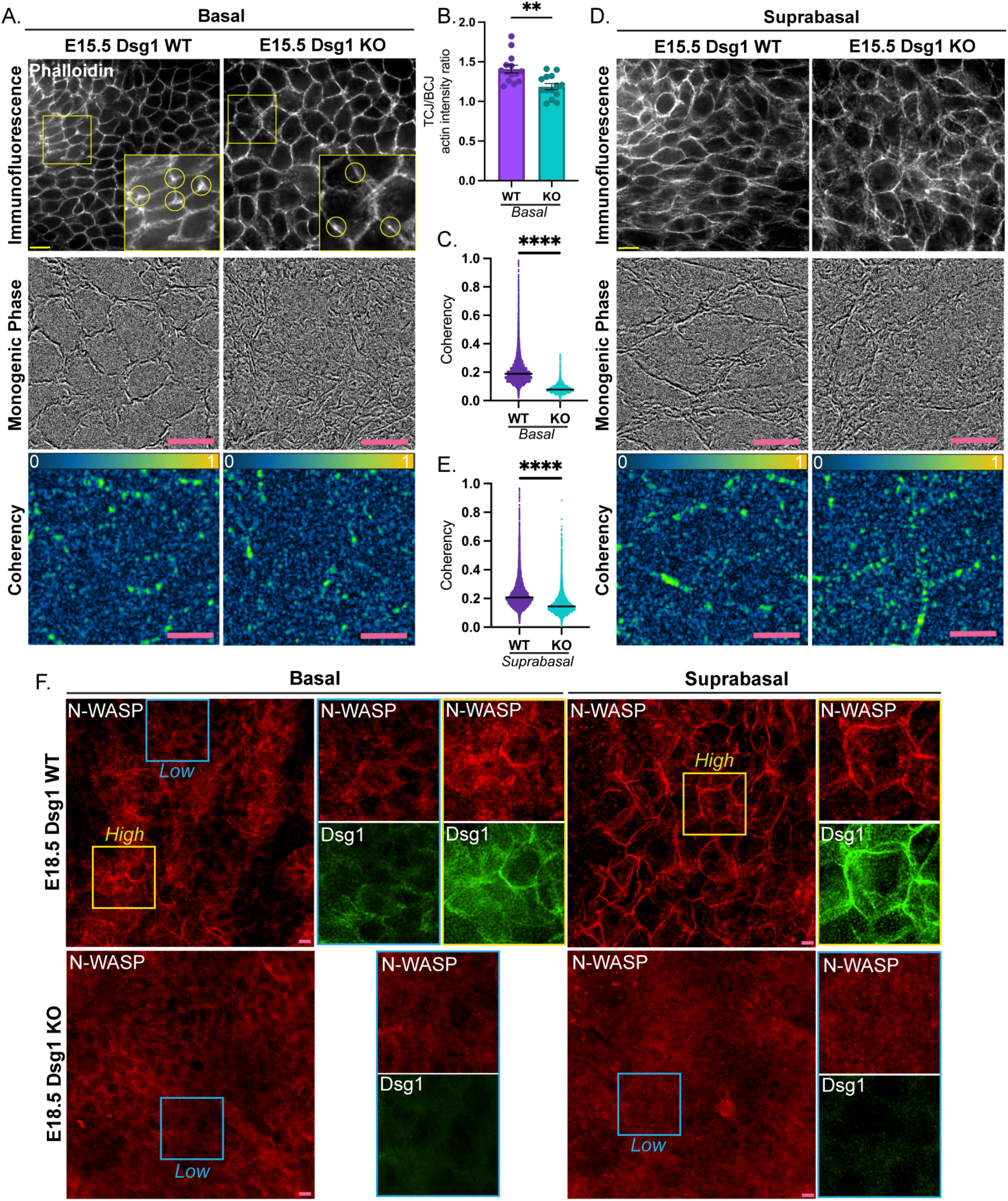
Dsg1 KO disrupts actin organization and the actin regulator, N-WASP, during early and late development, respectively. A) Representative basal layer images of E15.5 dorsal skin whole-mounts stained with phalloidin, comparing Dsg1 WT and Dsg1 KO. The upper panels show phalloidin immunofluorescence, with magnified insets highlighting phalloidin intensity at tricellular junctions (TCJs) relative to bicellular junctions (BCJs). The middle panels show monogenic phase analysis, and the lower panels show corresponding coherency maps, with coherency values represented on a scale from 0 to 1. B) Quantification of actin enrichment at TCJs, measured by phalloidin intensity and normalized to the average phalloidin intensity of the three adjacent BCJs, represented by yellow circles in zoom insets shown in (A, upper panels). C) Quantification of basal layer coherency shown in (A, lower panel). D) Representative suprabasal layer images of E15.5 dorsal skin whole-mounts stained with phalloidin, comparing Dsg1 WT and Dsg1 KO. The upper panels show phalloidin immunofluorescence, the middle panels show monogenic phase analysis, and the lower panels show corresponding coherency maps, with coherency values represented on a scale from 0 to 1. E) Quantification of suprabasal layer coherency shown in (B, lower panel). F) Representative images of E18.5 dorsal skin whole-mounts from Dsg1 WT and Dsg1 KO stained for N-WASP and Dsg1. The left panels show the basal layer, while the right panels show the suprabasal layer. Magnified insets highlight regions of low (blue box) and high (yellow box) Dsg1 expression and the corresponding N-WASP staining patterns across the basal and suprabasal layers. Data in (A,D) and (F) representative of n=2-3 and n=5-7 embryos per genotype, respectively. Individual data points in B,C and E represent single-cell measurements pooled from n = 2-3 embryos per genotype/timepoint; cell-level distributions were performed using the two-sample Kolmogorov-Smirnov test. Scale bar 20µm. p-values are represented by (**) = 0.01, (****) = 0.0001.

Our previous work showed that Dsg1 recruits the actin modulator, Arp3, to the plasma membrane, promoting actin remodeling that redistributes tension away from actin-anchoring AJs and facilitates cell delamination in vitro^10^. However, it remained unclear what links Dsg1 to the Arp2/3 machinery. N-WASP activates Arp2/3-dependent actin polymerization^38^ and is required for epidermal homeostasis, as keratinocyte-specific N-WASP deletion causes skin barrier defects and atopic dermatitis-like inflammation^39,40^. In E18.5 Dsg1 WT whole-mounts N-WASP strongly colocalized with Dsg1 in both the basal (Krt14+) and suprabasal (Krt10+) layers. In contrast, N-WASP failed to concentrate at cell-cell interfaces in E18.5 Dsg1 KO epidermis (**Figure 5F, Figure S9A-B**), suggesting that Dsg1 promotes junctional localization of N-WASP during epidermal development.

### Nck1 is a novel binding partner of Dsg1

The disruption of cortical actin organization and junctional N-WASP localization suggested that Dsg1 may couple membrane-associated complexes to actin assembly machinery. A publicly available desmoplakin (Dsp) BioID screen identified Nck1^41^, an SH2/SH3 protein^42^ that promotes Arp2/3-dependent actin assembly through interactions with N-WASP and cortactin, as a potential linker^43^. Sequence analysis identified a putative Nck1 binding motif (YSTV) at Y1042 in the Dsg1 C-terminal domain^44–46^ (**Figure 6A**). To test its role, we generated a Dsg1 Y1042F-GFP mutant predicted to disrupt phosphotyrosine-dependent Nck1 recruitment^42,43,46^(**Figure 6B**). To address whether Nck1 and Dsg1 exhibit close proximity that is disrupted by the Y1042F mutation, we utilized the proximity ligation assay (PLA) in reconstituted 3D neonatal human epidermal keratinocytes (NHEKs) epidermal raft equivalents transduced with GFP-tagged shCtrl, shDsg1, shDsg1 with Dsg1 rescue (Dsg1 KD + Dsg-GFP), and shDsg1 with the Dsg1-Y1042F mutant (Dsg1 KD + Dsg1-Y1042F). Significant PLA signals were detected in shCtrl and Dsg1-rescued rafts but not rafts with Dsg1-Y1042F expression (**Figure 6C-E**). These results suggest that the Dsg1-Nck1 interaction requires the Y1042 region and support a model in which Nck1 links Dsg1 to N-WASP:Arp3-dependent actin remodeling (**Figure 6F**).

**Figure 6:**
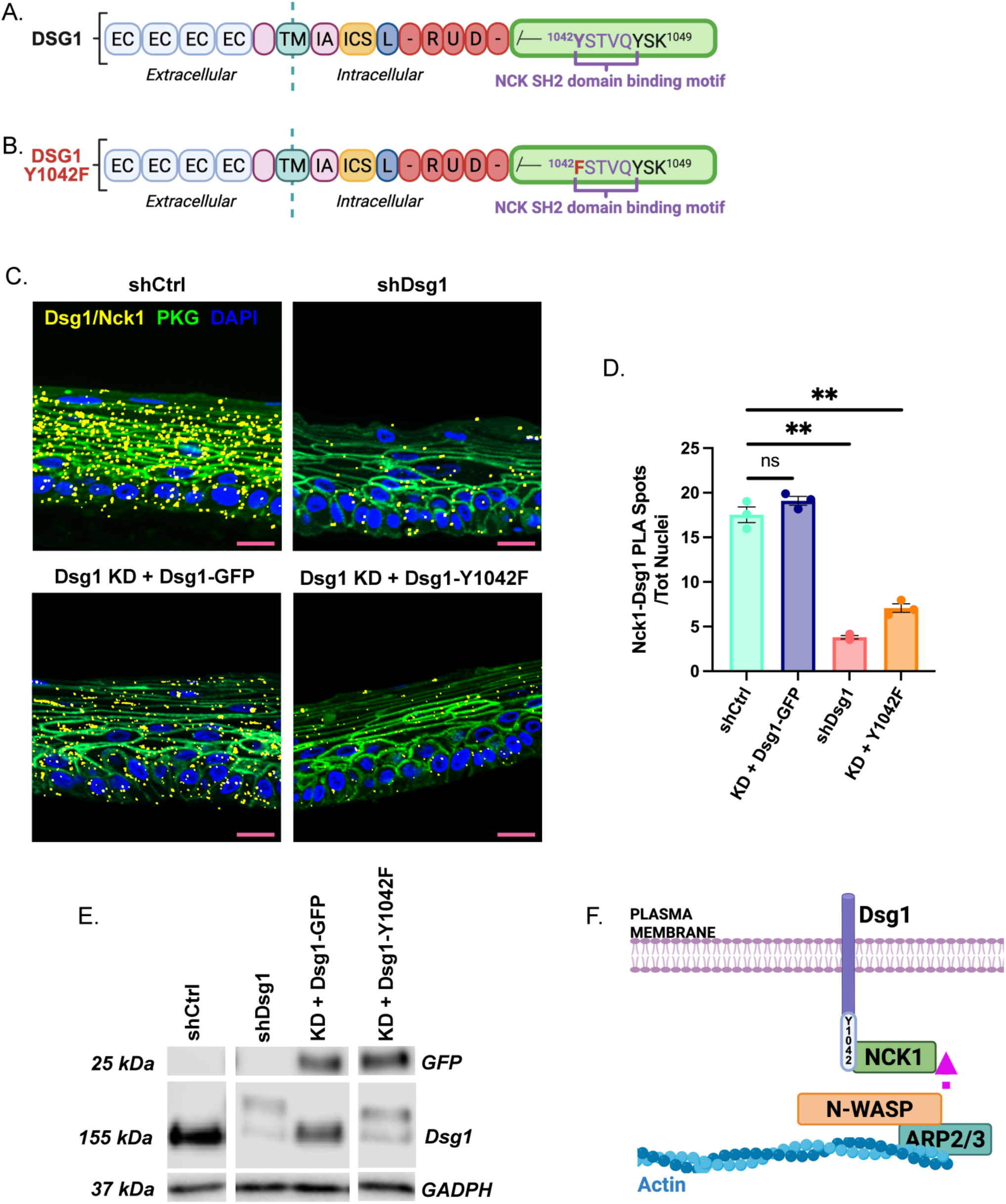
Nck1 is a novel binding partner of Dsg1, at the Dsg1 C-terminal domain sequence Y1042. A) Schematic of the Dsg1 protein sequence highlighting the C-terminal Nck1-binding motif (YSTVQ), which contains a tyrosine residue at Y1042. B) Schematic of the mutant Dsg1-Y1042F construct highlighting the Y1042F substitution (Y->F) within the C-terminal Nck1-binding motif. C) Representative images of proximity ligation assays (PLA) performed in Day 6 3D organotypic cultures generated from NHEKs transduced with shCtrl, shDsg1 (Dsg1 KD), Dsg1 KD + Dsg1-Y1042F (KD + Y1042F), or Dsg1 KD + Dsg1-GFP (KD + Dsg1-GFP) GFP-tagged constructs and stained for plakoglobin (PKG) and DAPI. Dsg1/Nck1 PLA puncta are shown in yellow. D) Quantification of Dsg1/Nck1 PLA puncta normalized to the total number of nuclei per 20X field, measured across 8-10 fields from each of three biological replicates, as shown in (C). E) Representative immunoblot confirming expression of the retroviral constructs used in (C). F) Schematic of the proposed interaction between Dsg1 and Nck1 at the Y1042-containing Nck1-binding motif, highlighting Nck1 as a potential link between Dsg1 and N-WASP-mediated Arp2/3-dependent actin polymerization. Statistical comparisons were performed using separate unpaired two-tailed Welch’s t-tests comparing each experimental condition with shCtrl. Scale bar 20µm. p-values are represented by (**) = 0.01; ns = not significant.

### The Dsg1-Nck1 interaction regulates actin remodeling required for proper stratification

Given our previous *in vitro* observation that Dsg1-dependent actin remodeling is associated with proper delamination, we tested the necessity of Nck1 as a bridge to actin polymerization machinery for stratification in a cell sorting assay. NHEKs expressing GFP/siCtrl or GFP/siNck1 were mixed 50:50 with non-transfected, unlabeled cells and allowed to stratify for 24 hours in high calcium media (HCM; 1.2mM). Parallel cultures maintained in low calcium media (LCM; 0.07mM) were used to establish the GFP-positive mixing ratio. Cells are stained for E-cadherin, and the transcription factor GRHL1, to mark cell membranes and stratified cells, respectively. In HCM, the number of GFP-positive/GRHL1-positive stratified cells were quantified relative to the total number of GFP-positive cells and the established mixing ratio (**Figure 7A**)^21^. Nck1 silencing resulted in a 4-fold reduction in stratification capacity compared to controls (**Figure 7B-D**). Similarly, N-WASP-silencing reduced stratification approximately 4-fold compared to controls (**Figure 7E-G**).

**Figure 7:**
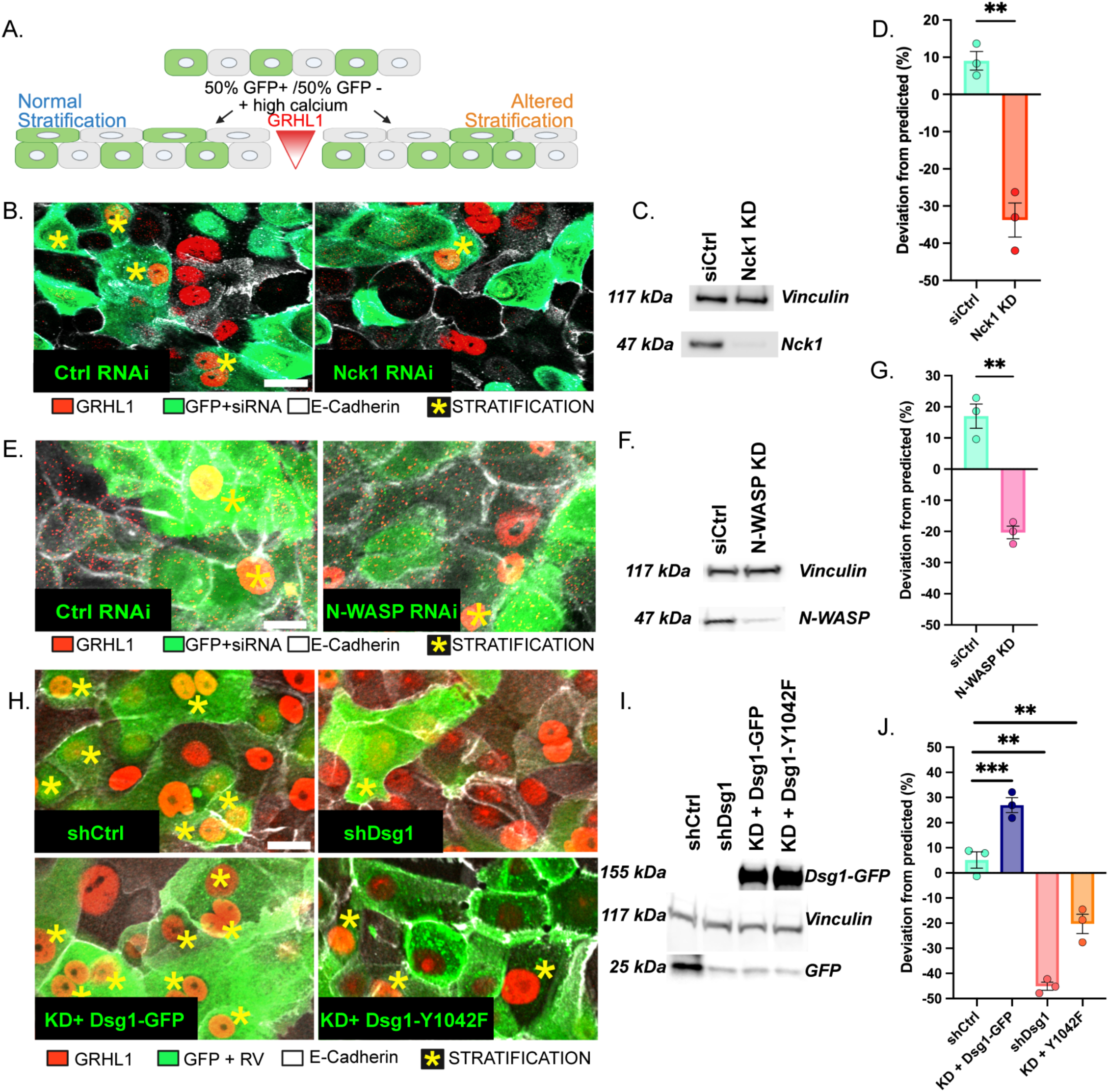
Dsg1-Nck1-N-WASP interaction is required for proper stratification. A) Schematic of the cell-sorting assay. B) Representative images of NHEKs transfected with siCtrl or Nck1 RNAi (Nck1 KD) and stained for E-cadherin, GFP, and GRHL1. Yellow asterisks indicate GFP+/GRHL1+ stratified cells used for quantification. C) Representative immunoblot confirming Nck1 knockdown, with vinculin shown as a loading control, for the conditions shown in (B). D) Quantification of GFP+/GRHL1+ stratified cells shown in (B). E) Representative images of NHEKs transfected with siCtrl or N-WASP RNAi (N-WASP KD) and stained for E-cadherin, GFP, and GRHL1. Yellow asterisks indicate GFP+/GRHL1+ stratified cells used for quantification. F) Representative immunoblot confirming N-WASP knockdown, with vinculin shown as a loading control, for the conditions shown in (E). G) Quantification of GFP+/GRHL1+ stratified cells shown in (E). H) Representative images of NHEKs transduced with shCtrl, shDsg1 (Dsg1 KD), Dsg1 KD + Dsg1-Y1042F (KD + Y1042F), or Dsg1 KD + Dsg1-GFP (KD + Dsg1-GFP) GFP-tagged constructs stained for E-cadherin, GFP, and GRHL1. Yellow asterisks indicate GFP+/GRHL1+ stratified cells used for quantification. I) Representative immunoblot confirming expression of the GFP-tagged constructs, with the vinculin as the loading control, for the conditions shown in (H). J) Quantification of GFP+/GRHL1+ stratified cells shown in (H). Data are presented as mean ± SEM from at least three biological replicates. Statistical comparisons were performed using unpaired two-tailed Welch’s t-tests for two-group comparisons or one-way ANOVA followed by Tukey’s multiple-comparisons test for experiments containing multiple conditions. Scale bars 10 µm. p-values are represented by (**) = 0.01, (***) = 0.001.

To test the role of the Dsg1 Y1042 Nck1 binding motif in stratification, NHEKs were transduced with GFP-tagged shCtrl, shDsg1, shDsg1 with Dsg1 rescue (Dsg1 KD + Dsg-GFP), and shDsg1 with Dsg1-Y1042F mutant (Dsg1 KD + Dsg1-Y1042F) and analyzed with the same stratification assay (**Figure 7A**). Dsg1 Y1042F expression resulted in a significant 3-fold decrease in stratification capacity, similar to the siNck1 and siN-WASP conditions. In contrast, Dsg1 rescue increased stratification 3-fold, consent with previous work^22^ (**Figure 7H-J**). Thus, disruption of the Y1042F Nck1-binding motif phenocopies loss of Nck1 or N-WASP, supporting a model in which Dsg1 recruits Nck1 to engage N-WASP-dependent actin remodeling required for proper stratification via delamination.

## Discussion

Using a new murine skin developmental atlas together with tissue- and cell-level analyses, we show that loss of Dsg1 alters the developmental programs underlying epidermal morphogenesis, while revealing tissue-level consequences and adaptations that emerge over time. Comparative scRNA-seq analysis identified altered cell-cycle and cytoskeletal programs in Dsg1 KO epidermis, pointing to changes in both proliferative and morphogenetic pathways. Importantly, these molecular changes were accompanied by a temporal defect in stratification, with reduced epidermal thickness at E14.5-E15.5 followed by recovery at later stages. The timing of this defect coincides with the emergence of Dsg1 and a developmental transition in stratification and terminal differentiation, processes consistent with established roles for Dsg1 in promoting differentiation and delamination in human *in vitro* models^19,20^. Thus, although epidermal thickness is restored by E18-P0^8^, our findings reveal an earlier requirement for Dsg1 in the timely progression of epidermal morphogenesis.

The enrichment of cell-cycle programs in Dsg1 KO keratinocytes was reflected by increased EdU incorporation, PCNA expression, and Rb phosphorylation during late development, particularly in transitional and suprabasal compartments. This persistent, spatially redistributed proliferation coincided with recovery of epidermal thickness and may represent an adaptive response to impaired early stratification. This connection may also be relevant to disease, as DSG1 haploinsufficiency causes palmoplantar keratoderma associated with epidermal hyperplasia^11^, while loss of desmoplakin, a cytoplasmic desmosome protein localized at the Dsg1 C-terminus, similarly enhances proliferation and G1-to-S progression^47^.

Dsg1 loss also produced temporally and compartmentally distinct changes in cell volume, shape, and actin organization. These changes span a recently described E15.5-E16.5 developmental transition during which evolving cell geometry and tissue mechanics create a barrier to delamination, requiring emerging mechanisms that enable basal cells to change shape and move suprabasally while maintaining tissue integrity^13,48^. Actomyosin remodeling is well positioned to meet this challenge, as dynamic changes in cortical tension allow epithelial cells to alter their geometry while maintaining adhesive contacts with neighboring cells^16^. Our findings that Dsg1 loss alters transitional cell geometry, actin coherency, and tricellular/bicellular junction enrichment ratio suggest that Dsg1 helps coordinate adhesive interactions with cytoskeletal remodeling needed to accommodate the changing mechanical environment of the developing epidermis. Dsg1-actin coupling may thereby facilitate the cell deformation and rearrangements required as stratification proceeds, consistent with our previous observation that Dsg1 contributes to the mechanical gradient across the epidermis^21^. Notably, epidermal thickness ultimately recovers despite persistent alterations in proliferative and morphogenetic programs, suggesting that adaptive mechanisms can compensate for the early consequences of Dsg1 loss.

To further investigate the mechanisms through which Dsg1 may regulate actin remodeling, we examined the distribution of N-WASP, an actin nucleation-promoting factor previously demonstrated to be essential for establishing the epidermal barrier^39,40^. N-WASP is also required for Arp2/3-dependent branched actin polymerization, which we previously demonstrated is initiated around desmosomes newly expressing Dsg1 ^10,44,49^. We observed that Dsg1 KO inhibits N-WASP association with the plasma membrane, resulting in a more diffuse cytoplasmic localization, suggesting that Dsg1 could influence the spatial regulation of N-WASP during epidermal differentiation. In this regard, N-WASP has also been shown to contribute to inflammation suppression through inhibition of IL-23, a cytokine we previously showed to be elevated and contribute to inflammation in Dsg1-deficient humans with SAM syndrome^8,40,50,51^.

Nck1 functions as an adaptor capable of coupling upstream signaling complexes to regulators of actin polymerization, including N-WASP, which promotes Arp2/3-dependent actin nucleation and branching^42,43^. Here, we determined that mutation of the C-terminal Y1042 residue is important for Nck1:Dsg1 co-localization at the membrane and that loss of Nck1, N-WASP and a Nck1-binding deficient mutant significantly reduced stratification capacity *in vitro*. Previous studies identified another SH2/3 family member, CrkL, as being a critical contributor to epidermal integrity^41^. While we cannot rule out a contribution of Crk binding to Dsg1 to epidermal development, its presence was not sufficient for maintaining stratification competence of Nck1-silenced keratinocytes in our experiments. These findings support a model in which Dsg1 helps spatially coordinate Nck1:N-WASP-dependent actin-regulatory machinery at cell-cell interfaces, thereby coupling intercellular adhesion to cytoskeletal remodeling during epidermal morphogenesis.

Collectively, our work fills gaps in the current publicly available embryonic atlases spanning from early (E14.5, E15.5) to late (E16.5, E18.5) embryonic stages of murine epidermal stratification and reveals how Dsg1 coordinates proliferative and morphogenetic programs during formation of the multilayered epidermis. Loss of Dsg1 delays normal stratification but is followed by proliferative and morphological programs that ultimately restore epidermal thickness, highlighting the plasticity of the developing epidermis. Mechanistically, we identify Nck1 and N-WASP as a link between the Dsg1 desmosomal complex and actin remodeling machinery required for proper stratification. These findings support a model in which differentiation-dependent expression of Dsg1 coordinates cell-cycle regulation with cytoskeletal remodeling to shape epidermal architecture and barrier formation, while revealing mechanisms that preserve tissue formation when this coordination is disrupted.

## Materials and Methods

### Animals

All procedures related to the housing, care, and utilization of embryos adhered to the Northwestern University Institutional Animal Care and Use Committee Protocols (ID IS00021648). C57BL/6 mice were housed in a barrier facility with controlled temperature, subjected to a 12-hour light cycle, and provided unrestricted access to food and water. The procedure for generating the Dsg1 knockout mouse (Dsg1 KO) was previously documented^8^. Mice were paired overnight and separated the following morning to facilitate timed pregnancies and embryos were collected on day E14.5, E15.5, E16.5, E18.5. Whole embryos or dorsal skin from both wild-type and Dsg1 KO embryos were collected.

### Primary Normal Human Keratinocytes

Primary keratinocytes were isolated in accordance with established protocols from human neonatal foreskin sourced from Northwestern University’s Skin Biology and Disease Resource-based Center (IRB Protocol #STU00009443)^52^. The undifferentiated keratinocytes were cultured in M154 media (ThermoFisher), supplemented with 0.07mM CaCl^2^, human keratinocyte growth supplement (HKGS), gentamicin, and amphotericin B. To induce differentiation in 2D cultures, the calcium concentration of the growth medium was elevated to 1.2mM CaCl^2^. Additionally, 3D organotypic epidermal raft cultures using primary keratinocytes were generated following established protocols^52^.

### scRNA Sample and Library Preparation

Single cells were isolated from embryonic dorsal skin harvested at E14.5, E15.5, E16.5, and E18.5 timepoints. Dorsal skin samples were first treated with dispase for 15min at 37°C, followed by treatment with Collagenase IV for 30 minutes at 37°C. After collagenase IV treatment, the tissue was extensively triturated and pelleted. The pelleted cells were resuspended with pre-warmed, fresh trypsin/EDTA and incubated for 5 min at 37°C. After incubation, cell suspensions were placed on ice and treated with 3% chelexed FBS before triturating. Cells were then passed through a 40µm filter, pelleted, and resuspended in PBS. Cells from two pups were combined for each sample per genotype and timepoint and from the combined suspension, 8,000-10,000 cells were used to generate scRNAseq libraries following manufacturer protocols (10x Genomics Chromium Single Cell v3.1 kit). The libraries were quality checked using an Agilent Bioanalyzer 2100 and DNA High Sensitivity kit (Agilent cat #5067-4626) before being sent out for sequencing and demultiplexing (Novogene Corp).

### Single-cell RNA-sequence Quality Control and Preprocessing

Single-cell RNA-sequencing data were processed using R 4.6.1 (R Core Team, 2021) and Seurat package (v5)^26^. Cells were filtered on a per-sample basis to remove low-quality or outlier cells, excluding cells with fewer than 500 detected genes or more than 10,000 detected genes, as well as cells with greater than 20% mitochondrial transcript content. Doublets were identified and removed using DoubletFinder (R 4.6.1, v2.0.6)^53^. Quality control covariates were calculated for each cell, including the percentage of mitochondrial genes (mt), histone genes (H[1-4]), hemoglobin genes (Hba, Hbb), ribosomal protein genes (Rps, Rpl), and Xist expression. Cell cycle scores were computed using canonical S-phase and G2/M-phase gene sets (cc.genes.updated.2019), and a cell cycle difference score (S.Score-G2M.Score) was calculated for downstream regression. For the construction of the wild-type (WT) reference atlas, data were log-normalized using NormalizeData, and 2,000 highly variable genes were identified using the variance-stabilizing transformation. Data were scaled using ScaleData, regressing out cell cycle effects and the above quality control covariates. For subclustering analyses, variable feature selection and regression parameters were adjusted as appropriate for the subsetted populations. Cell cycle effects were regressed during data scaling for the purpose of dimensionality reduction and clustering, to prevent cell cycle–driven variance from dominating low-dimensional embeddings. Importantly, cell cycle scores were retained as metadata and were not removed from the expression matrix. These scores were subsequently used for downstream analyses to assess genotype-dependent changes in proliferative state, enabling evaluation of altered cell cycle dynamics independently of their influence on clustering structure.

### Dimensionality Reduction, Batch Correction, and Clustering

Principal component analysis (PCA) was performed on the variable genes, and batch effects across samples were corrected using Harmony^54^, grouping by “orig.ident”. A UMAP embedding was generated from the Harmony-corrected space using a selected subset of Harmony dimensions (dims = 1, 4, 8, 9, 11, 14, 15), chosen based on inspection of variance contribution and biological interpretability. Nearest-neighbor graphs and clustering were performed in Harmony space, and clusters were annotated using established marker genes to define keratinocyte subpopulations.

### Reference-based Label Transfer and Projection of the Dsg1 KO dataset

To evaluate transcriptional changes induced by gene knockout while preserving the structure of the Dsg1 WT atlas, a reference-based projection strategy was employed in R 4.6.1. Dsg1 WT cells were used as the reference dataset, and Dsg1 knockout (Dsg1 KO) cells were treated as the query. Dsg1 KO cells were preprocessed using the same normalization, quality control, cell cycle scoring, regression, and PCA workflow as the Dsg1 WT reference. Transfer anchors were identified using FindTransferAnchors, leveraging the Harmony-corrected Dsg1 WT embedding as the reference reduction. Dsg1 KO cells were then projected into the Dsg1 WT low-dimensional space using MapQuery, which transferred Dsg1 WT cluster annotations and generated projected UMAP coordinates for Dsg1 KO cells. This approach enabled direct comparison of Dsg1 KO cells within the Dsg1 WT-defined transcriptional landscape without recomputing a joint embedding. Label transfer and merged reference-query objects were used exclusively for visualization and annotation alignment, but not for statistical comparisons, to avoid biases introduced by projection-based embeddings.

### Downstream Analyses

Quantitative gene expression analyses were performed on the jointly processed Dsg1 WT and Dsg1 KO dataset to ensure expression values reflected observed transcript abundance in each condition. Differential expression analysis was performed using Seurat’s v5 FindMarkers function (Wilcoxon rank-sum test), with genes considered differentially expressed at an adjusted p-value < 0.05 and an average log2 fold-change > 0.25. Functional enrichment analysis was conducted using Metascape^27^, focusing on Gene Ontology (GO) pathways and curated pathway databases, with significance defined as p < 0.05 with input lists were limited to 3,000 genes.

### Antibodies and Reagents

Primary antibodies for immunofluorescence and/or immunoblotting were anti-E-cadherin (BD Bioscience, BD61081), anti-Keratin-14 (Biolegend, Poly906), anti-Keratin-10 (Biolegend, Poly1905), anti-Keratin-10 (Progen, GP-K10), anti-PCNA (Cell Signaling, 2585), anti-N-WASP (Cell Signaling, 4848S), anti-Nck (Abcam, ab32120), P124 (anti-Dsg1 extracellular domain, Progen, 651111), 27B2 (anti-Dsg1 cytodomain, Life Technologies, 32600), 4B2 (anti-Dsg1 C-terminal domain^55^), anti-GRHL1 (ThermoFisher Scientific, PA531734), anti-GFP (Aves labs, GFP-1020), JL-8 (anti-GFP, Clontech), anti-vinculin (clone hVIN-1, V9131, Sigma-Alrich), pRb (Cell Signaling, 8516), and anti-Plakoglobin (1408, Aves Laboratories). Secondary antibodies for immunoblotting were goat-anti-mouse, goat-anti-rabbit, and goat-anti-chicken (LICOR). Secondary antibodies for immunofluorescence were goat-anti-mouse (IgG and IgG2a), goat-anti-rabbit, goat-anti-chicken, and goat-anti-guinea pig linked to fluorophores 488, 568, or 647(Alexa Fluor, Life Technologies). Phallodin 647 (Invitrogen, A22287 or A30107) was used to label F-actin, DAPI (Sigma-Alrich) was used to stain nuclei.

### Whole-Mount Tissue Staining and Image Analysis

Embryos collected from E15.5, E16.5, and E18.5 timed litters were fixed in 4% PFA in PBS with calcium and magnesium (Corning) for 45min-2hrs at room temperature while rocking. After fixation, the dorsal skin was carefully dissected from the embryos using micro-dissecting scissors, non-serrated forceps, and pipettes tips affixed with a single toothbrush bristle^56^. The pelt was cut into sections and transferred to a 24-well plate and incubated in a blocking/permeabilizing solution (5% normal goat serum (NGS), 0.5% Triton-X 100) rocking overnight at 4°C. The tissues were rinsed several times in PBS and incubated in a cocktail of primary antibodies diluted in 5% normal goat serum, 0.25% Triton-X 100 + overnight at 4°C. Following primary incubation, samples were rinsed several times in PBS and incubated with secondary antibodies diluted in 5% NGS, 0.25% Triton and incubated overnight at 4°C. After secondary incubation, tissues were washed in PBS and mounted epidermis side down in a 24-well glass bottom dish using polyvinyl alcohol. A coverslip was placed on top of the tissue and gently pressed down to keep the tissue flat for imaging. Whole-mounts images were acquired using a Nikon AXR confocal microscope with a Plan Apo λD 20× OFN25 DIC N2 objective or a Nikon CSU-W1 SoRa with a Plan Apo Lambda λS 40×C Sil objective.

### *In utero* EdU Incorporation

On the day of collection, pregnant dams received an intraperitoneal injection of 10mg/kg EdU (5-Ethynyl-2-deoxyuridine, Invitrogen, E10187) in their lower abdomen. Dams were euthanized after 1 hour and immediately dissected to remove the embryos. All embryos were fixed for 48hrs in 10% neutral buffered formalin and processed for paraffin embedding.

### Immunofluorescence, Histology, and Image Acquisition

Paraffin-embedded tissue sections from mouse dorsal skin, whole embryos, and 3D epidermal raft cultures were baked at 60°C overnight. Deparaffinization of tissue sections was achieved using xylene/ethanol, and 0.5% Triton X-100 was utilized for tissue permeabilization. Antigen retrieval was performed by heating samples to 95°C in a 1.8mM citric acid and 8.2mM sodium citrate buffer at pH 6.0. Sections were blocked in 1% BSA with 2% normal goat serum for 1 h at 37°C and incubated with primary antibodies overnight at 4°C in a humid chamber and secondary antibodies at 37°C for 1 h followed by mounting with ProLong Gold antifade reagent. Samples undergoing EdU detection were incubated for an additional hour at room temperature following blocking but before incubation with primary antibodies using the Invitrogen Click-iT EdU Cell Proliferation Kit (C10339). Images were acquired with an epifluorescence microscope (Axio Imager Z2, Carl Zeiss) equipped with an Apotome.2 slide module, Axiocam 503 Mono digital camera, X-Cite Mini+ System, and a Plan-Apochromat 20x/0.8 M27 objective.

Primary keratinocytes were cultured on glass-bottom 24 well dishes. Fixation was carried out using either a 4% paraformaldehyde solution for 10-15 minutes at room temperature. For paraformaldehyde-fixed samples, 0.2% Triton X-100 was employed to permeabilize the cells for 10 minutes at room temperature. Subsequently, cells underwent blocking with a 1% BSA solution for 30 minutes at room temperature. Monolayers were incubated for 30 minutes at room temperature in blocking solution (1% BSA with 2% normal goat serum, or 1% BSA alone if a primary goat antibody was used). Primary and secondary antibodies were diluted in the blocking solution and applied to cells for either 1-2 hours at 37°C or overnight at 4°C. Images were acquired with an epifluorescence microscope (Axio Imager Z2, Carl Zeiss) equipped with an Apotome.2 slide module, Axiocam 503 Mono digital camera, X-Cite 120 LED Boost System, and Plan-Apochromat 20x/0.8 M27 objective.

### Proximity Ligation Assay

For the proximity ligation assay (PLA) tissue samples were blocked and incubated with primary antibodies as described above. Following primary antibody incubation, the PLA protocol for 2D and tissue samples was performed according to manufacturer’s instructions. Images were acquired with an epifluorescence microscope (Axio Imager Z2, Carl Zeiss) equipped with an Apotome.2 slide module, Axiocam 503 Mono digital camera, X-Cite 120 LED Boost System, and Plan-Apochromat 20x/0.8 M27 objective objective. Additional information on troubleshooting, image acquisition, and analysis^57^.

### FIJI Analyses

#### Dorsal Skin Tissue Cross Section Analysis

Epidermal thickness was quantified using 8-10 line scans across 8-10 acquired images, using H&E images or E-cadherin staining to mark the living layers and keratin-14 to denote epidermis from the dermal compartment. PCNA and EdU staining was manually quantified using ImageJ on maximum intensity projections (MIP) of 8-10 acquired images. A ratio of EdU or PCNA positive nuclei to total nuclei per 8-10 image 20x fields were manually obtained using the cell counter plug in. Only interfollicular cells were included in the assessment. Additionally, PCNA positive cells that were positive for keratin-14 and/or keratin-10 were manually counted. PLA spots were counted in ImageJ using the automated protocol described^57^. The apical-to-basal width ratio of wedge-shaped cells was manually quantified by identifying keratin-14/keratin-10 double-positive cells and using E-cadherin staining to delineate and measure their apical and basal widths to calculate basal/apical width ratio. The quantification of each image field per embryo was averaged and then used to run statistics using Prism software (version 10.0). For each timepoint comparison, Welch’s T-Test was used to generate statistics.

#### Dorsal Skin Whole-Mount Analysis

##### Volume Calculation

Images were analyzed using a combination of Fiji and Imaris software. Calculation of cell volumes was done assuming a frustum of a pyramid, using the following formula:

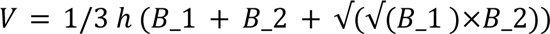

Where B1 and B2 are the areas of the top and bottom layers of each cell, and h is the height of the cell. Results of clearly defined cells were confirmed by generating iso-surfaces in Imaris and automatically calculating the cell volume. Not all cells were defined enough for segmentation (despite pre-processing steps) for automatic rendering, due to penetration depth or confluency within the tissue, therefore all volumes were obtained as previously described. The aspect ratio was calculated as the proportion of width and height dimensions of the cell.

#### Coherency and Monogenic Phase Analysis

Coherency and monogenic phase analyses were performed using Multiscale Directional Analysis from MonogenicJ (OrientationJ plugin in Fiji). We specifically obtained Laplace monogenic components, structure tensor features including orientation and coherency, and monogenic analysis including wavenumber (local spatial frequency), modulus (local strength of the structural feature), phase (local symmetry type), and directional Hilbert (a 90 degree phase-shifted signal along the dominant structure path. The parameters used for this analysis were Nb of scale 2, and Sigma [Tensor] 3. Coherency maps were processed using a Green Fire Blue LUT, to highlight coherency.

#### Tricellular Junction MFI Ratio Calculation

The skeletonize 2D/3D Fiji plugin was used to calculate the MFI ratio of tricellular junctions (TCJ) to the value of bicellular junctional (BCJ) lines. Images were background corrected, and masks were generated to retain only the cell borders. Every vertex point and their edges were identified by skeletonizing the various fields of view. The average intensity of the 3 BCJ lines linked to any individual TCJ was calculated, and the ratio obtained by dividing the TCJ intensity over the BCJ average.

### Immunoblotting

Keratinocytes were lysed using Urea Sample Buffer (8M deionized urea, 1% SDS, 60 mM Tris, pH 6.8, 10% glycerol, 0.1% pyronin-Y, 5% beta-mercaptoethanol). Proteins were separated using SDS-PAGE electrophoresis before transfer to a nitrocellulose membrane (0.2 or 0.45 mm pore size). Samples were resolved on 7.5% SDS/PAGE gels purchased from BioRad and subjected to 80-100 volts in Tris/Glycine buffer containing 0.1%SDS. Membranes were blocked with either 5% Milk or LICOR Intercept (PBS) Blocking Buffer, (LI-COR). Primary and secondary antibodies were diluted in the blocking buffer. Immunoreactive proteins were visualized using NIR fluorescence or chemiluminescence using the Odyssey FC Imaging system (LI-COR).

### siRNA-mediated Transfection and siRNAs

Keratinocytes were transiently transfected using Amaxa Nucleofector System (Lonza) for electroporation of cells according to manufacturer’s instructions with siRNA. Following trypsinization, 5×10^5^-2×10^6^ keratinocytes were pelleted and resuspended in 100µl Ingenio Electroporation Solution with 5µl siRNA per 1×10^6^ keratinocytes (20µM stock concentration and electroporated using program X-001. Dharmacon siGENOME SMARTpool siRNA’s (Horizon Discovery) were used to target human N-WASP (WASL, M-006444-02-005) and human Nck1 (M-006354-01-005) compared to non-targeting siRNA pool#2 (D-001206-14).

### Plasmids and Viral Transduction

pLZRS constructs (shCtrl-GFP, shDsg1-GFP, Dsg1-GFP, and Dsg1-Y1042F-GFP) were transiently transfected into Phoenix cells (provided by G. Nolan, Stanford University, Stanford, CA) using Lipofectamine 2000. Phoenix cells successfully transfected with the plasmid were selected using 1µg/ml puromycin. Following a 24-48hr incubation at 32°C, retrovirus-containing Phoenix cell supernatant was collected and concentrated using Amicon Ultra-15 Centrifugal Filter Units (EMD Millipore). Keratinocytes were infected with concentrated retrovirus diluted in growth medium and 4µg/ml polybrene at 20-40% confluency for 1-1.5 hours at 32°C.

### Stratification Assay

Stratification assays and analysis were performed as previously described for Dsg1-GFP stratification studies allowing GFP labeled keratinocytes mixed with unlabeled keratinocytes to stratify for 24hrs^21^. For Nck1 and N-WASP depletion, Nck1 and N-WASP siRNA were transfected into undifferentiated GFP expressing keratinocytes before mixing with non-transfected and unlabeled keratinocytes in a known ratio. For Nck1-binding deficient stratification experiments, GFP-labeled keratinocytes expressing shCtrl, shDsg1, shDsg1+WT-Dsg1, and shDsg1+Y1042F-Dsg1 were transduced into undifferentiated keratinocytes before mixing with non-transduced and unlabeled keratinocytes in a known ratio. Following 24hrs of stratification, 4% paraformaldehyde-fixed cells were processed as described above and stained for E-cadherin, GRHL1, GFP and DAPI. Confocal images with 0.5mm z-stacks were obtained to calculate the ratio of stratified cells that are GFP positive. The percentage of GFP-positive stratified cells per condition was compared to the predicted percentage of cells that express GFP after mixing with unlabeled cells to obtain the percent deviation from predicted.

### Quantification and Statistical Analysis

A minimum of three independent experiments (biological replicates) were conducted for each experiment. For 2D experiments, keratinocytes isolated from distinct foreskin samples were treated as biological replicates. In immunofluorescence studies, a minimum of eight 20x fields from three embryos across each timepoint were used to quantify EdU, PCNA, and pRb data normalized to total number of nuclei, a minimum of eight 20x fields from three 3D organotypic rafts were used to quantify PLA plots normalized to total number of nuclei, and 2D cell sorting assay encompassed a minimum of 100 cells per 20x field. All images within each experiment were captured using consistent parameters. FIJI/ImageJ software (NIH) was utilized for the analysis of fluorescence intensity, area, perimeter, and volume used for cell shape index calculation^34^. Graphing and statistical analyses were conducted using R (4.6.1) or GraphPad Prism (version 10.0) software, with specific tests indicated in the figure legends. A p-value less than 0.05 was deemed statistically significant. The graphs depict the mean of the independent experiments, represented by large circles, along with individual data points colored according to each independent experiment, including paired samples when applicable. The error bars denote the standard error of the mean (SEM).

## Acknowledgements

Funding: The authors thank the invaluable input from members of the Green lab, and conversations with Drs. Lisa Manning, Sara Wickström, and other colleagues in the cell adhesion and epithelial differentiation communities. This work was supported by National Institutes of Health grants R01 AR041836, R01 AR043380 and CA228196 to Kathleen Green and F31 AR086611 to Brieanna Jarrell and R01 HD107841, R01 AR081103, R01 AG100048 to Rui Yi. Imaging work was performed at the Northwestern University Center for Advanced Microscopy (RRID: SCR_020996) generously supported by NCI CCSG P30 CA060553 awarded to the Robert H Lurie Comprehensive Cancer Center. Super resolution and confocal microscopy was performed on a Nikon SoRa system, purchased through the support of NIH 1S10OD032270-01. We thank the STEM Core of the Northwestern University Skin Biology and Diseases Resource-based Center (P30 AR075049) for human keratinocytes used in this manuscript. **Author contributions:** Conceptualization: B.L.J., R.D., M.H., L.G., and K.J.G. Methodology: B.L.J., R.D., M.H., M.D.N., J.L.K., L.G., D.W., M.R., C.N., Z.R., and K.J.G. Investigation: B.L.J., R.D., M.H., M.D.N, X.T., R.H., L.G., and J.L.K. Visualization: B.L.J., R.D., and Z.R. Supervision: L.G., Z.R., and K.J.G. Writing-original draft: B.L.J, M.H. and K.J.G. Writing-review and editing: C.N., M.R., L.G., R.H., and M.H.. Data, code, and materials availability: The sequencing data and codes will be available via GEO and github once the paper is accepted for publication. The rest of data needed to evaluate and reproduce the results in the paper are present in the paper and/or the Supplementary Materials.

## Supplemental Figure Legends

**Figure S1:**
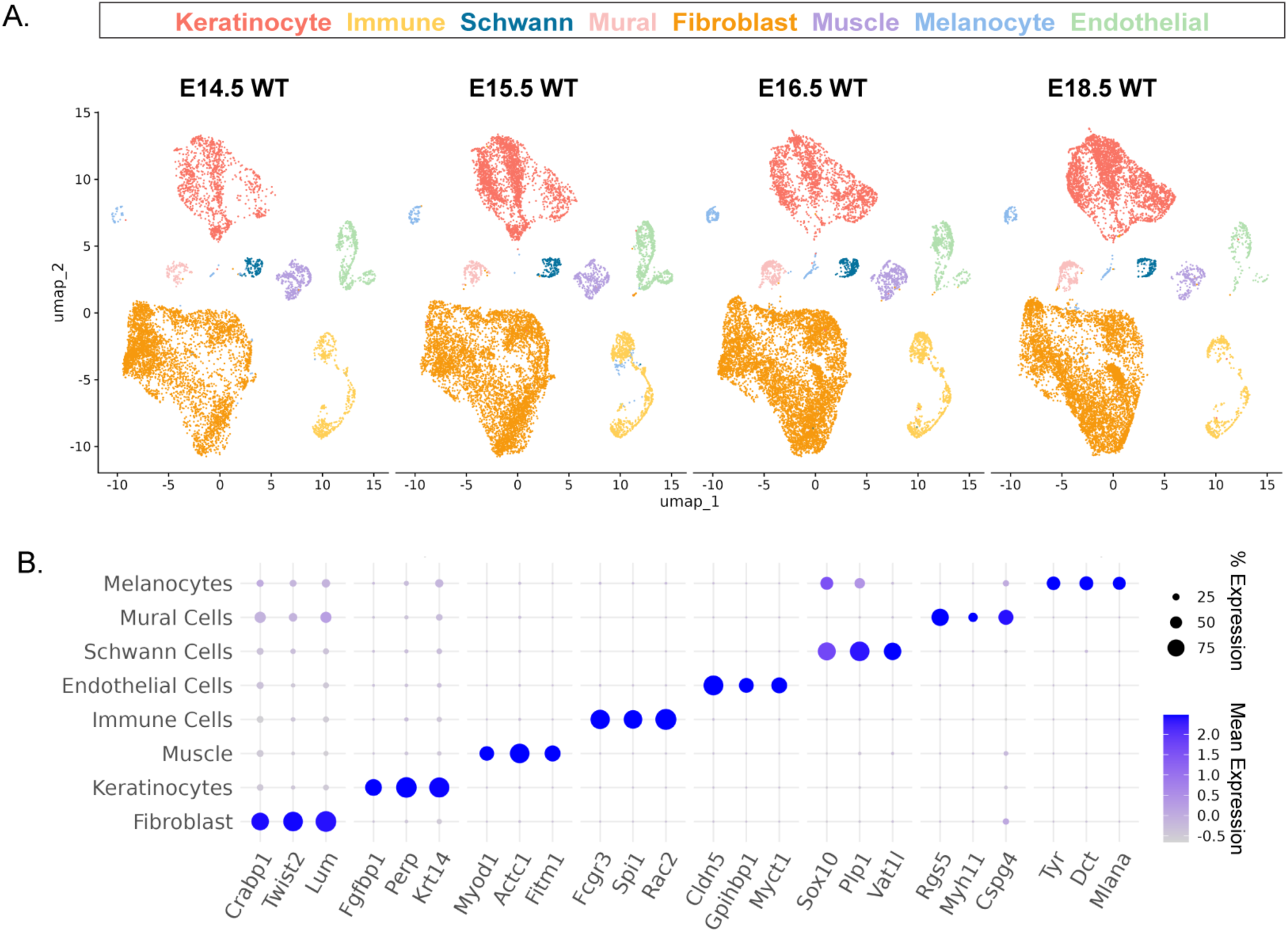
Dsg1 WT scRNA-seq murine skin atlas identifies eight epidermal and dermal cell populations across development. A) Dsg1 WT UMAPs at each developmental timepoint (E14.5, E15.5, E16.5, and E18.5) showing eight major epidermal and dermal cell populations resolved across development. B) Dot plot of canonical marker gene expression used to identify and annotate the eight cell populations shown in (A).

**Figure S2:**
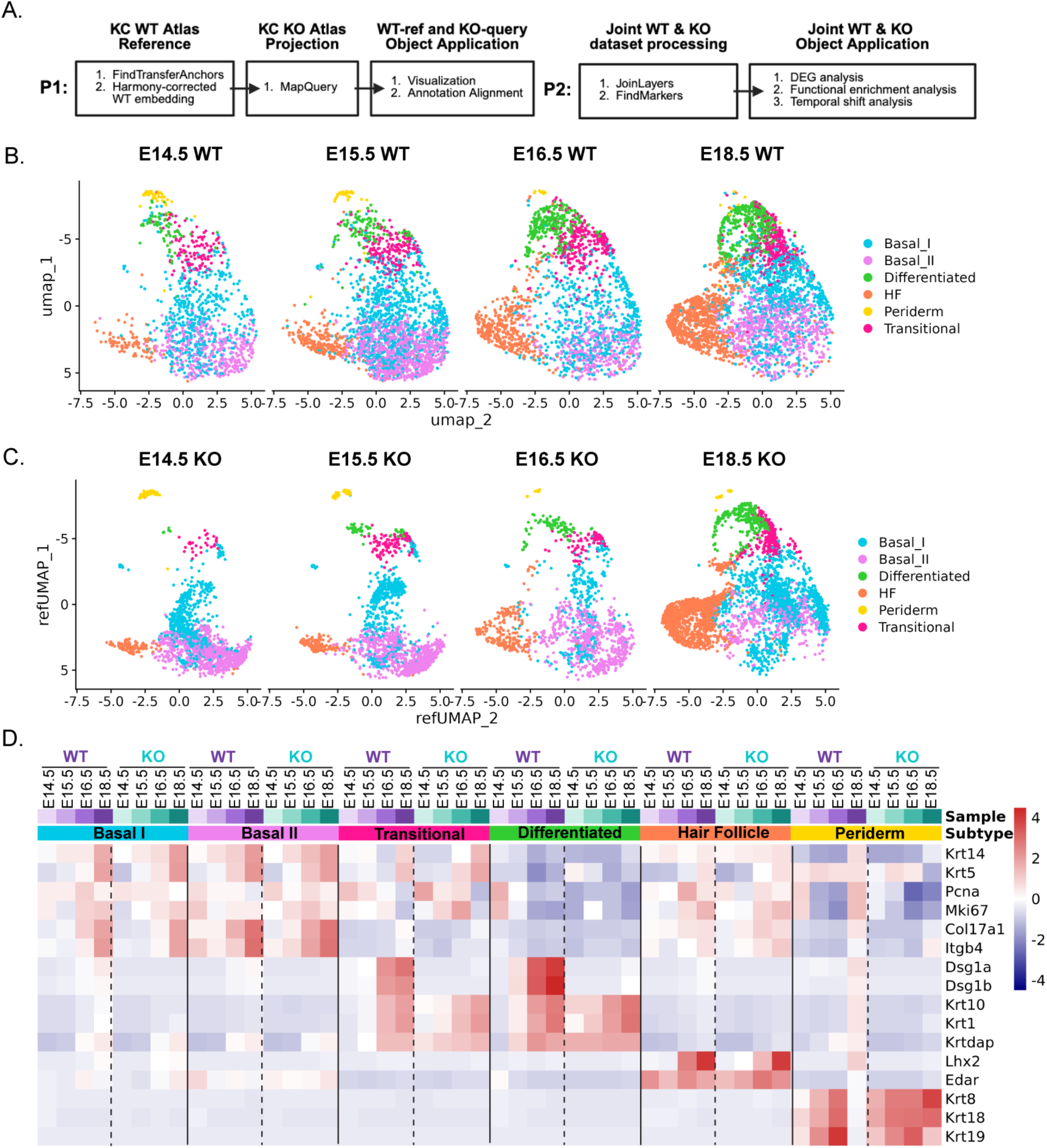
Workflow of keratinocyte-specific subclustering to generate keratinocyte WT-reference object and joint WT-KO object for downstream analysis. A) Workflow for generation of the Dsg1 WT keratinocyte reference atlas and projection of the Dsg1 KO query onto the WT reference, alongside preprocessing and integration of Dsg1 WT and KO datasets to generate a joint WT+KO object for downstream differential gene expression (DEG) analysis. B) Dsg1 WT reference UMAPs at each developmental timepoint (E14.5, E15.5, E16.5, and E18.5) showing six keratinocyte subpopulations resolved across epidermal development. C) Dsg1 KO query reference-mapped UMAPs (refUMAPs) at each developmental timepoint (E14.5, E15.5, E16.5, and E18.5), showing the six keratinocyte subpopulations projected onto the corresponding Dsg1 WT reference. D) Heatmap of Dsg1 WT reference and Dsg1 KO query cells showing marker gene expression used to annotate the six keratinocyte subpopulations (Basal I, Basal II, Transitional, Differentiated, Hair Follicle, and Periderm) across developmental timepoints E14.5, E15.5, E16.5, and E18.5.

**Figure S3:**
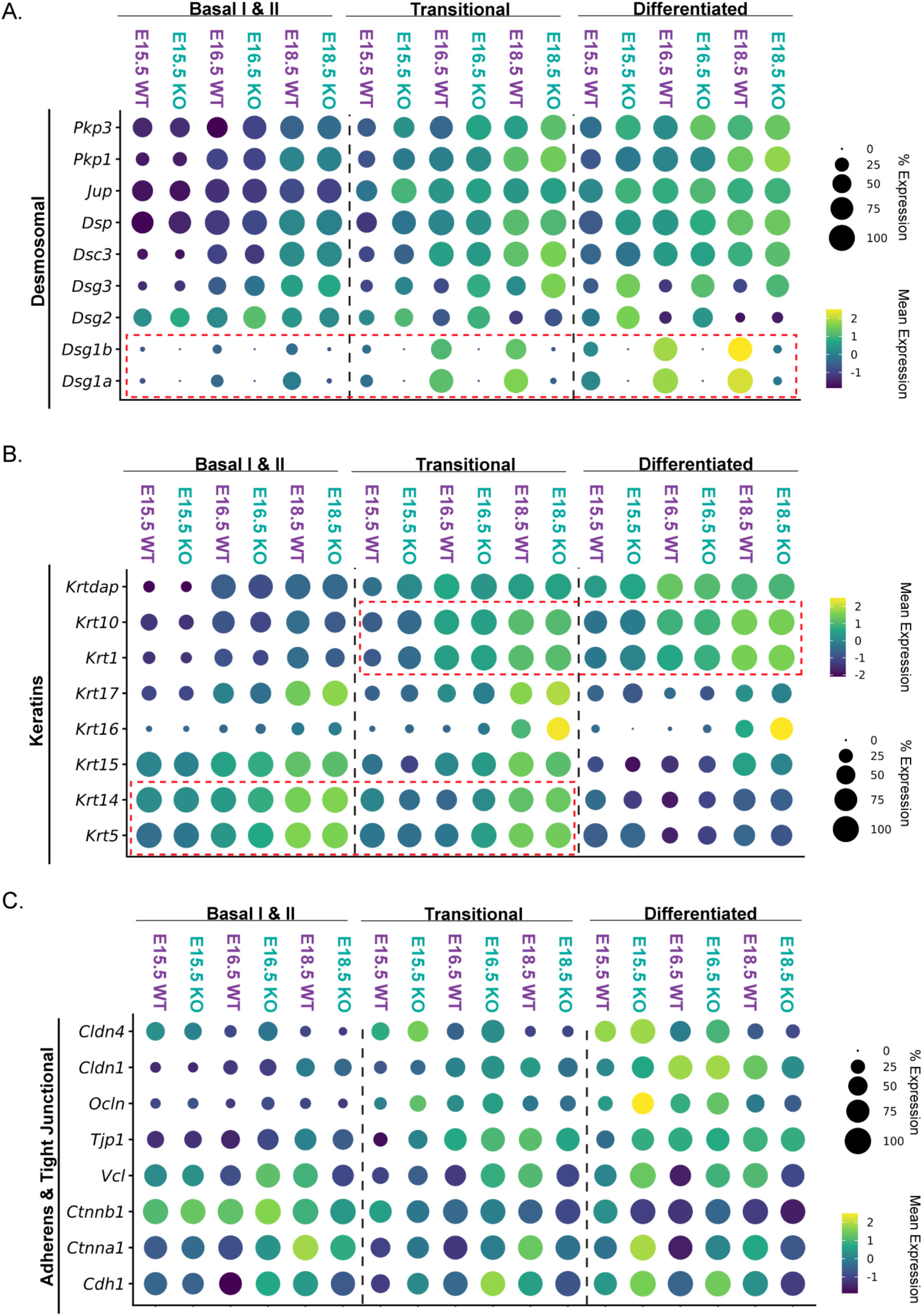
Dsg1 WT and Dsg1 KO keratinocyte subpopulations exhibit distinct epidermal keratin and cell junction gene expression profiles across development. A) Dot plot of selected desmosomal-related genes across E15.5, E16.5, & E18.5 in combined Basal I & II, Transitional, and Differentiated subpopulations, comparing Dsg1 WT and Dsg1 KO. Note the gradual increased expression of *Dsg1a/b* from Basal I & II to Transitional to Differentiated subpopulations across all Dsg1 WT timepoints (red dotted box). B) Dot plot of selected epidermal keratin genes across E15.5, E16.5, & E18.5 in combined Basal I & II, Transitional, and Differentiated subpopulations, comparing Dsg1 WT and Dsg1 KO. Note the increased expression of *Krt5/Krt14* and *Krt1/10* in the Transitional subpopulation (red dotted boxes). C) Dot plot of selected Adherens Junctional-related genes (*Cdh1, Ctnna1, Ctnnb1, Vcl*) and Tight Junctional-related genes (*Tjp1, Ocln, Cldn1, Cldn4*) across E15.5, E16.5, & E18.5 in combined Basal I & II, Transitional, and Differentiated subpopulations, comparing Dsg1 WT and Dsg1 KO.

**Figure S4:**
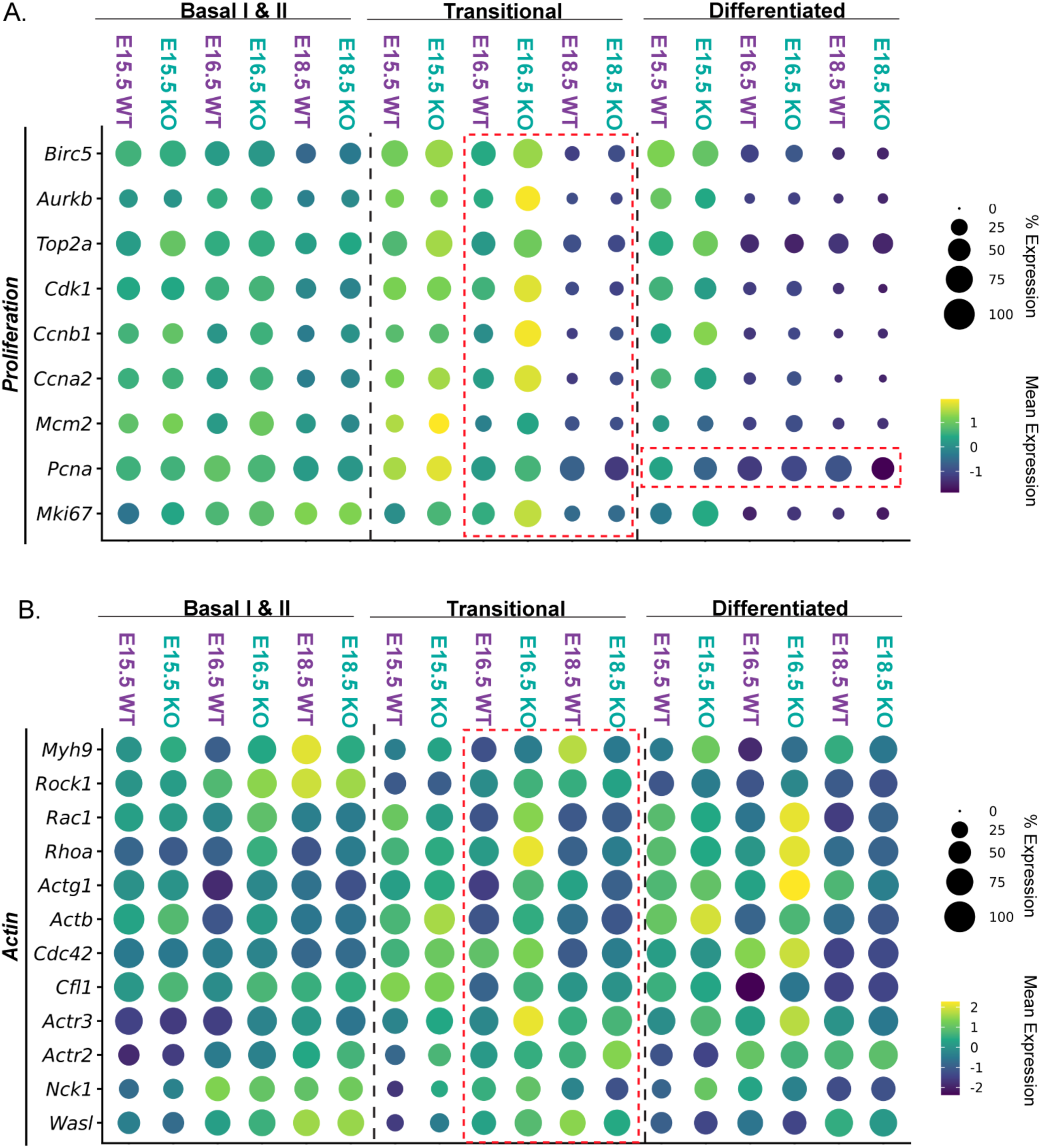
Dsg1 WT and Dsg1 KO keratinocyte subpopulations exhibit distinct cell-cycle and actin-associated gene expression profiles across development. A) Dot plot of selected cell proliferation-related genes across E15.5, E16.5, and E18.5 in combined Basal I & II, Transitional, and Differentiated keratinocyte subpopulations, comparing Dsg1 WT and Dsg1 KO. Note the increased expression of proliferation-related genes in Dsg1 KO Transitional and Differentiated subpopulations compared with Dsg1 WT at E16.5 and E18.5 (large red dotted box). Additionally, note the changes in *PCNA* expression within the Differentiated subpopulation across developmental timepoints (small red dotted box). B) Dot plot of selected actin-associated genes across E15.5, E16.5, and E18.5 in combined Basal I & II, Transitional, and Differentiated keratinocyte subpopulations, comparing Dsg1 WT and Dsg1 KO. Note the increased expression of actin-associated genes in Dsg1 KO Transitional and Differentiated subpopulations compared with Dsg1 WT at E16.5 and E18.5 (red dotted box).

**Figure S5:**
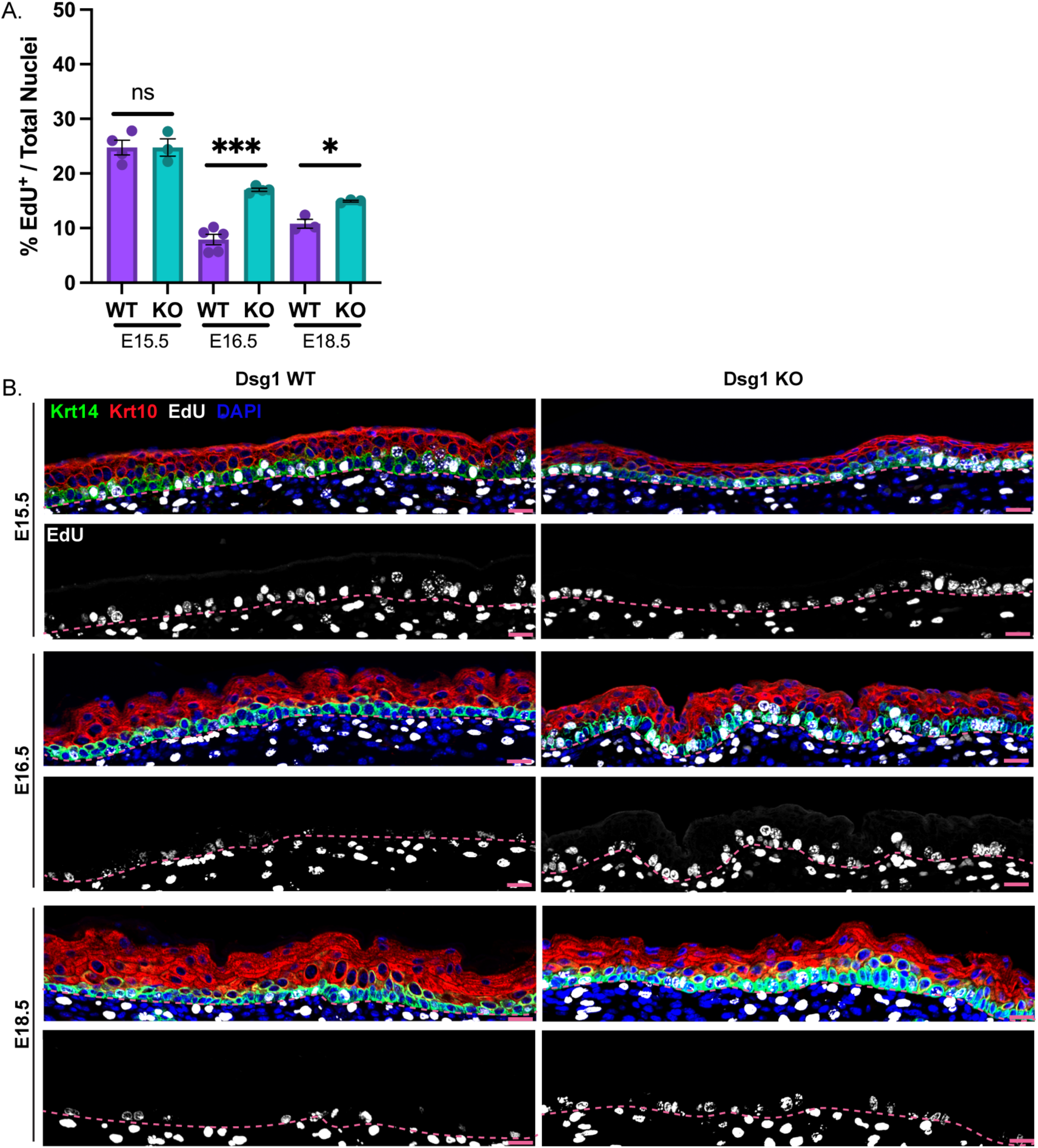
Dsg1 KO results in increased EdU incorporation during later epidermal development. A) Quantification of the percentage of EdU-positive cells detected by EdU Click-iT chemistry across developmental timepoints E15.5, E16.5, and E18.5, comparing Dsg1 WT and Dsg1 KO. B) Representative images of dorsal skin tissue sections from Dsg1 WT and Dsg1 KO at E15.5, E16.5, and E18.5 stained for Keratin-14 (Krt14), Keratin-10 (Krt10), EdU, and DAPI, corresponding to the quantification shown in (A). Data are presented as mean ± SEM from at least three biological replicates. Statistical comparisons were performed using separate unpaired two-tailed Welch’s t-tests comparing Dsg1 WT and Dsg1 KO within each developmental timepoint in (A). Scale bars 20 µm. p-values are represented by (*) = 0.05, (***) = 0.001; ns = not significant.

**Figure S6:**
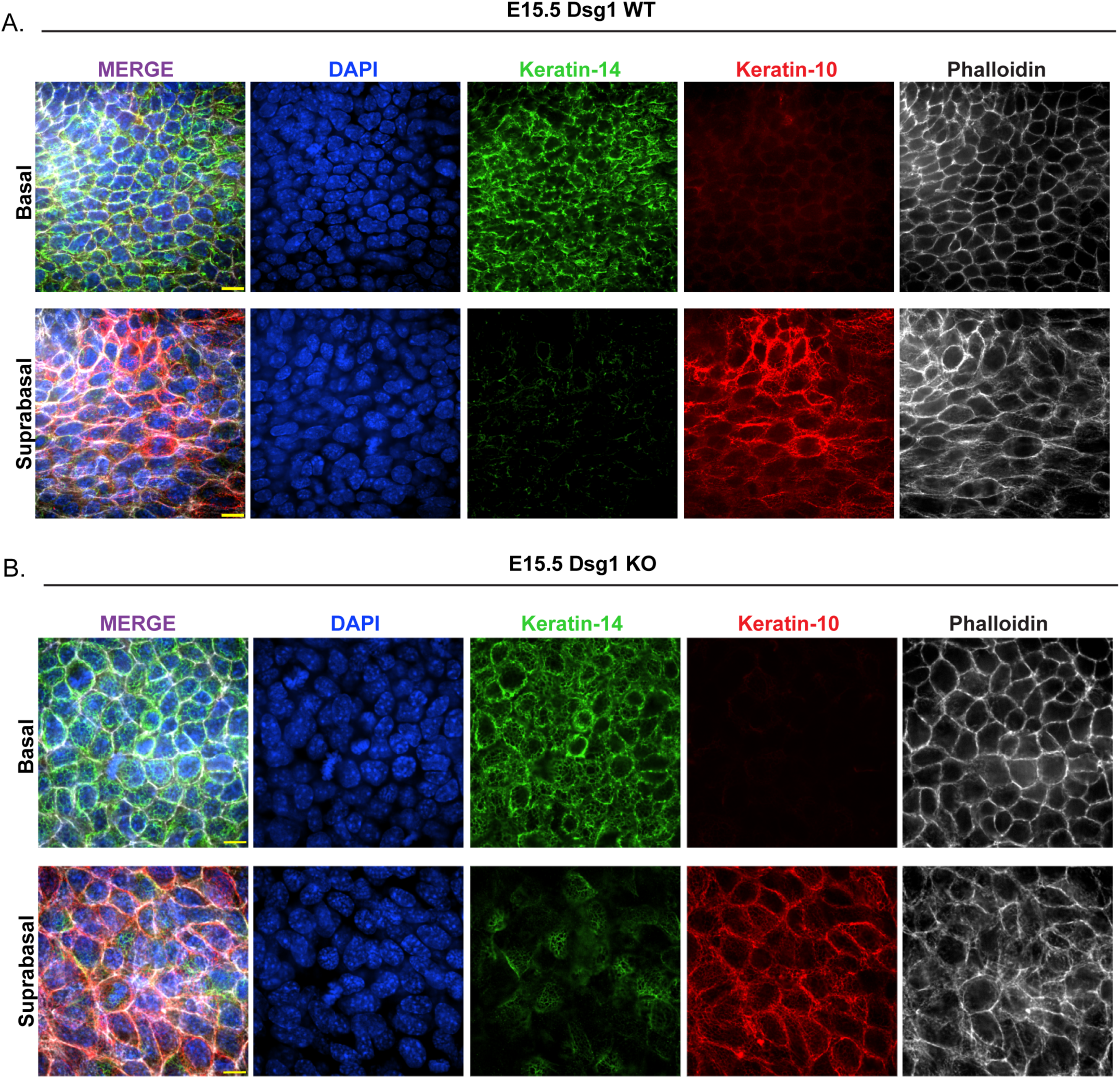
Whole-mount staining demonstrates layer-specific keratin expression at E15.5. A-B) Representative images of E15.5 Dsg1 WT and Dsg1 KO dorsal skin whole-mounts stained for Keratin-14, Keratin-10, and DAPI, with phalloidin staining to visualize cortical actin at cell boundaries. Keratin-14 and Keratin-10 staining was used to identify the basal (Krt14+) and suprabasal (Krt10+) epidermal layers, respectively. Scale bars 20 µm. Images representative of n=2-3 embryos per genotype.

**Figure S7:**
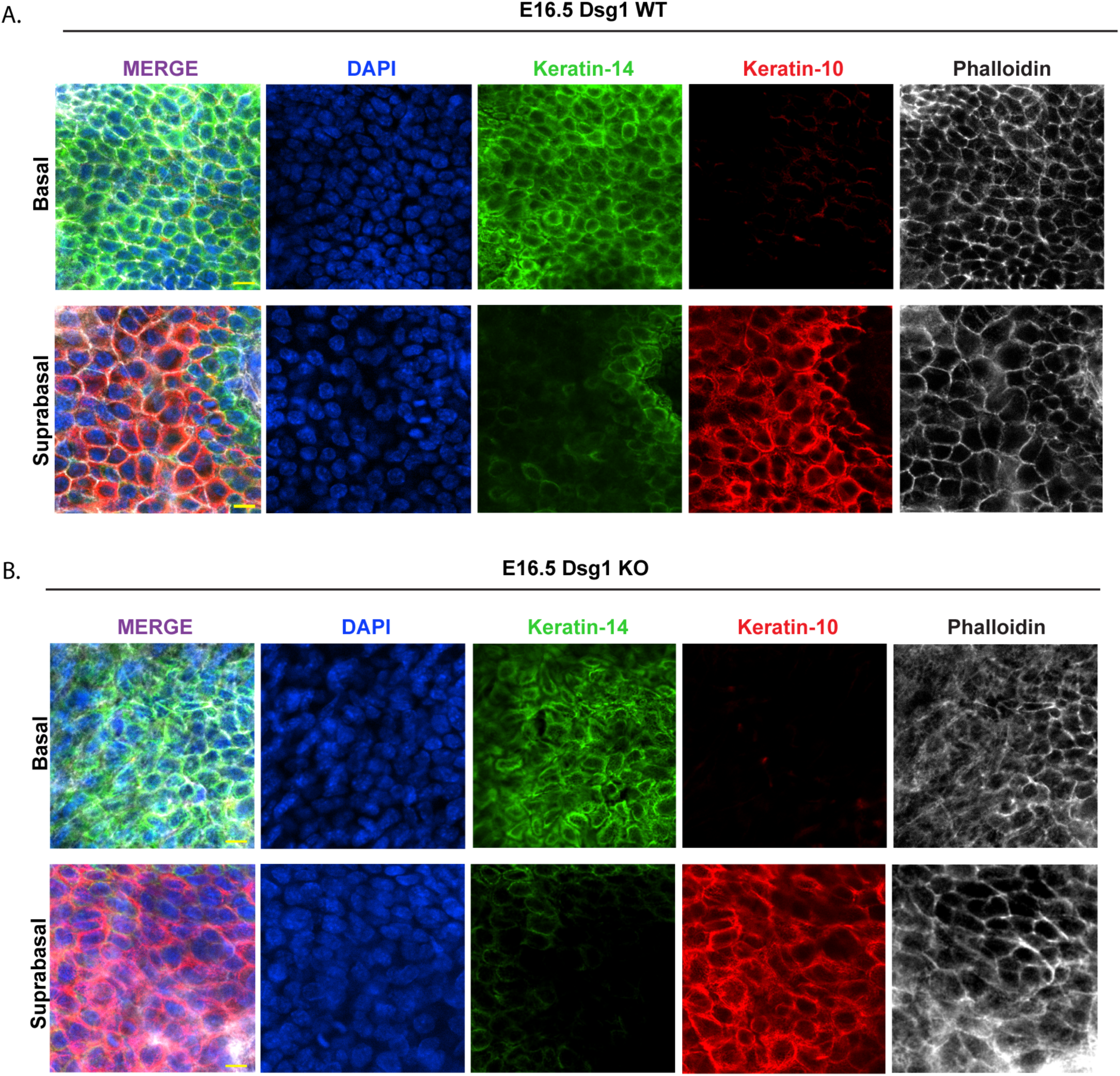
Whole-mount staining demonstrates layer-specific keratin expression at E16.5. A-B) Representative images of E16.5 Dsg1 WT and Dsg1 KO dorsal skin whole-mounts stained for Keratin-14, Keratin-10, and DAPI, with phalloidin staining to visualize cortical actin at cell boundaries. Keratin-14 and Keratin-10 staining was used to identify the basal (Krt14+) and suprabasal (Krt10+) epidermal layers, respectively. Scale bars 20 µm. Images representative of n=3 embryos per genotype.

**Figure S8.**
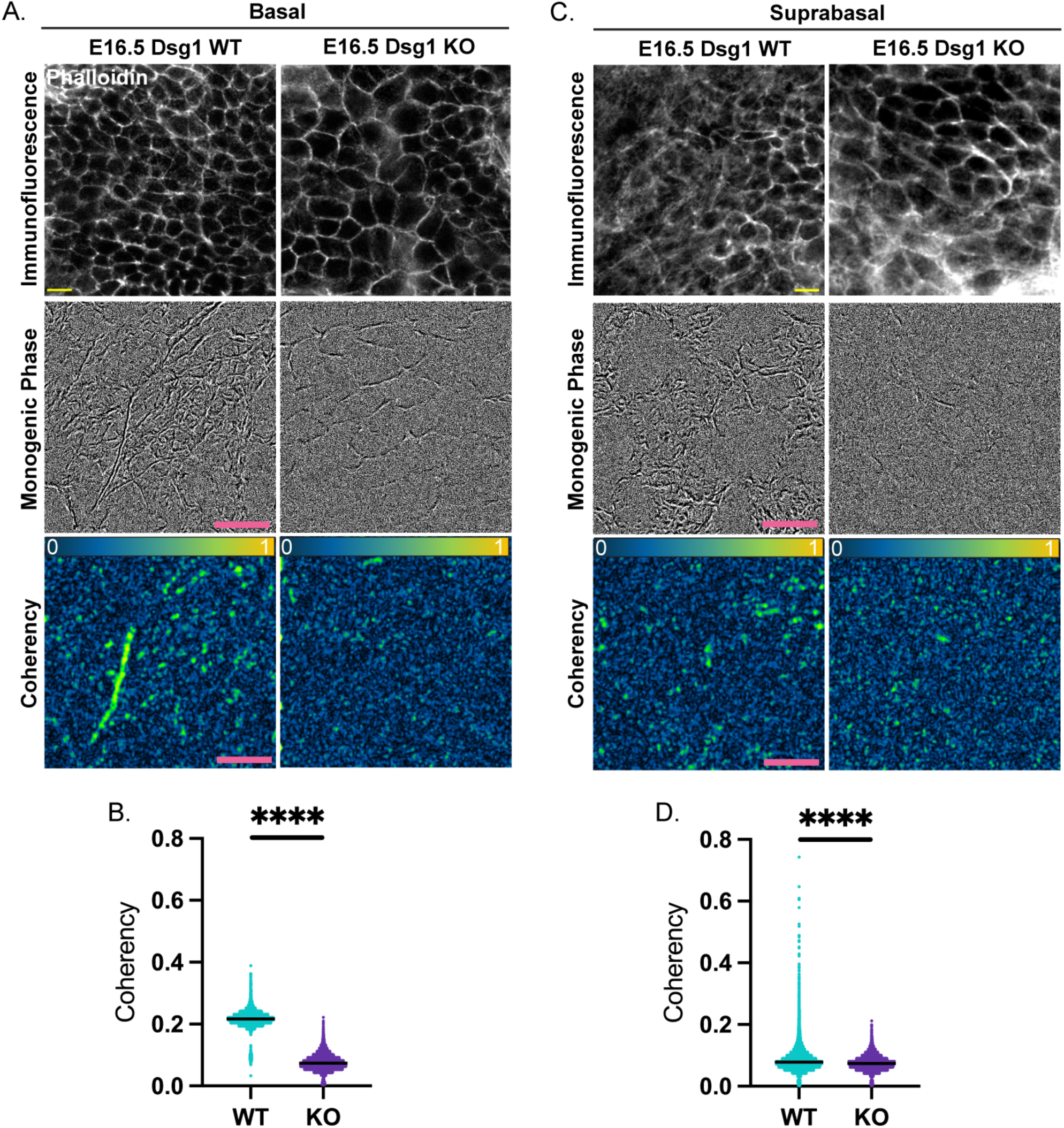
Dsg1 KO disrupts actin organization at E16.5, consistent with alterations observed at E15.5 (*shown in* Figure 5). A) Representative basal layer images of E16.5 dorsal skin whole-mounts stained with phalloidin, comparing Dsg1 WT and Dsg1 KO. The upper panels show phalloidin immunofluorescence, the middle panels show monogenic phase analysis, and the lower panels show corresponding coherency maps, with coherency values represented on a scale from 0 to 1. B) Quantification of basal layer actin coherency shown in (A, lower panels). C) Representative suprabasal layer images of E16.5 dorsal skin whole-mounts stained with phalloidin, comparing Dsg1 WT and Dsg1 KO. The upper panels show phalloidin immunofluorescence, the middle panels show monogenic phase analysis, and the lower panels show corresponding coherency maps, with coherency values represented on a scale from 0 to 1. D) Quantification of suprabasal layer actin coherency shown in (D, lower panels). (B,D) Individual data points represent single-cell measurements pooled from n = 3 embryos per genotype/timepoint; cell-level distributions were performed using the two-sample Kolmogorov-Smirnov test. Scale bar 20µm. p-values are represented by (****) = 0.0001.

**Figure S9.**
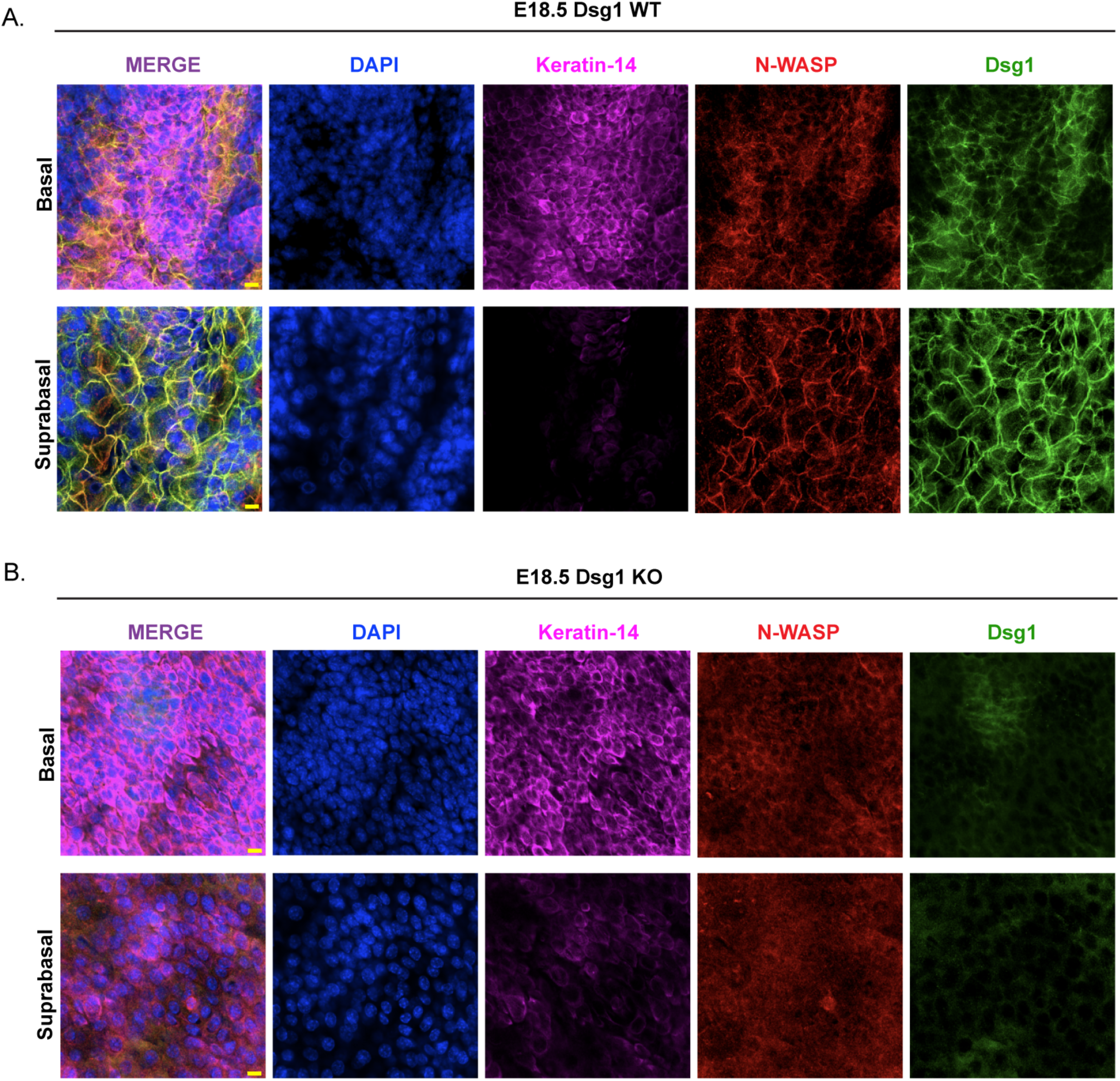
Whole-mount staining demonstrates layer-specific keratin expression at E18.5. A-B) Representative images of E18.5 Dsg1 WT and Dsg1 KO dorsal skin whole-mounts stained for Keratin-14, N-WASP, Dsg1, and DAPI. Keratin-14 staining was used to distinguish the Krt14+ basal layer from the overlying Krt10+ suprabasal layer for analysis of N-WASP and Dsg1 localization. Scale bars 20 µm. Images representative of n=5-7 embryos per genotype.

